# The polyadenylase PAPI is required for virulence plasmid maintenance in pathogenic bacteria

**DOI:** 10.1101/2024.10.11.617751

**Authors:** Katherine Schubert, Micah Braly, Jessica Zhang, Michele E. Muscolo, Hanh N. Lam, Karen Hug, Henry Moore, Joshua W. McCausland, Derfel Terciano, Todd Lowe, Cammie F. Lesser, Christine Jacobs-Wagner, Helen Wang, Victoria Auerbuch

## Abstract

Many species of pathogenic bacteria harbor critical plasmid-encoded virulence factors, and yet the regulation of plasmid replication is often poorly understood despite playing a critical role in plasmid-encoded gene expression. Human pathogenic *Yersinia*, including the plague agent *Y. pestis* and its close relative *Y. pseudotuberculosis*, require the type III secretion system (T3SS) virulence factor to subvert host defense mechanisms and colonize host tissues. The *Yersinia* T3SS is encoded on the IncFII <u>p</u>lasmid for *<u>Y</u>ersinia* <u>v</u>irulence (pYV). Several layers of gene regulation enables a large increase in expression of *Yersinia* T3SS genes at mammalian body temperature. Surprisingly, T3SS expression is also controlled at the level of gene dosage. The number of pYV molecules relative to the number of chromosomes per cell, referred to as plasmid copy number, increases with temperature. The ability to increase and maintain elevated pYV plasmid copy number, and therefore T3SS gene dosage, at 37°C is important for *Yersinia* virulence. In addition, pYV is highly stable in *Yersinia* at all temperatures, despite being dispensable for growth outside the host. Yet how *Yersinia* reinforces elevated plasmid replication and plasmid stability remains unclear. In this study, we show that the chromosomal gene *pcnB* encoding the polyadenylase PAP I is required for regulation of pYV plasmid copy number (PCN), maintenance of pYV in the bacterial population outside the host, robust T3SS activity, and *Yersinia* virulence in a mouse infection model. Likewise, *pcnB*/PAP I is also required for robust expression of the *Shigella flexneri* virulence plasmid-encoded T3SS. Furthermore, *Yersinia* and *Shigella pcnB*/PAP I is required for maintaining normal PCN of model antimicrobial resistance (AMR) plasmids whose replication is regulated by sRNA, thereby increasing antibiotic resistance by ten-fold. These data suggest that *pcnB*/PAP I contributes to the spread and stabilization of virulence and AMR plasmids in bacterial pathogens, and is essential in maintaining the gene dosage required to mediate plasmid-encoded traits. Importantly *pcnB*/PAP I has been bioinformatically identified in many species of bacteria despite being studied in only a few species to date. Our work highlights the potential importance of *pcnB*/PAP I in antibiotic resistance, and shows for the first time that *pcnB*/PAP I reinforces PCN and virulence plasmid stability in natural pathogenic hosts with a direct impact on bacterial virulence.

**Author Summary:** Many pathogens carry extrachromosomal DNA elements known as plasmids, which encode genes that confer bacterial virulence or antimicrobial resistance (AMR). Acquisition of these plasmids by bacteria can lead to the emergence of new pathogenic traits and the spread of AMR, yet the mechanisms by which plasmids are retained in bacterial populations particularly in the absence of selective pressure remain incompletely understood. Here we show that the major bacterial polyadenylase enzyme PAP I, encoded by the *pcnB* gene, is critical for the human pathogen *Yersinia pseudotuberculosis* to maintain its native virulence plasmid as well as AMR plasmids. Very little is known about the process of polyadenylation in bacteria, or the post-transcriptional addition of adenosine residues to the 3’ end of transcripts. This study represents the first demonstration that PAP I-mediated polyadenylation contributes to bacterial pathogenesis.

## Introduction

Many species of pathogenic bacteria harbor plasmids that encode genes important for virulence or antimicrobial resistance (AMR). Virulence plasmids are typically characterized by their relatively large size (>40kb) and low plasmid copy number (PCN), which refers to the average number of plasmid copies per cell or chromosome equivalent [1]. In addition to virulence or AMR genes, these plasmids typically contain additional genes important for plasmid replication, PCN control, and plasmid maintenance. Virulence plasmids are commonly found in members of the Enterobacteriaceae family, which includes several species of Gram-negative pathogens. For example, *Yersinia pseudotuberculosis* and the related plague agent *Yersinia pestis* naturally harbor a ∼70 kb low copy number virulence plasmid known as the *<u>p</u>lasmid for <u>Y</u>ersinia <u>v</u>irulence* (pYV), also called pCD1 in *Y. pestis* [2–4]. Given that *Y. pestis* has killed over 200 million people since antiquity [5], pYV has played a grim but important role in human history.

The pYV plasmid encodes several factors important for *Yersinia* virulence, including a major virulence factor known as the type III secretion system (T3SS) [6–8]. The T3SS is used to inject a series of effector proteins known as *<u>Y</u>ersinia* <u>o</u>uter <u>p</u>roteins (Yops) directly into the cytosol of target host cells during infection. Yops dampen the host immune response and promote *Yersinia* pathogenesis, making pYV essential for *Yersinia* virulence [3, 9–13]. Expression and activity of the T3SS exerts a metabolic burden on the bacteria. Therefore, *Yersinia* repress expression of the T3SS at low environmental temperatures when the T3SS is dispensable, but induce T3SS expression at mammalian body temperature, 37°C [14–19]. Several mechanisms contribute to this temperature dependency. For example, the master regulator of the T3SS is only translated at 37°C due to control by an RNA thermometer [18, 20]. More recently, it was reported that temperature-dependent changes in T3SS gene expression is also regulated at the level of gene dosage via changes in pYV PCN [21]. At environmental temperatures such as 26°C, pYV PCN has been reported to be low (∼1-2 copies per chromosome). Upon exposure to mammalian temperatures of 37°C, pYV PCN has been shown to be rapidly upregulated to 3+ copies per chromosome. Importantly, this ability to upregulate pYV PCN and T3SS gene dosage is essential for normal T3SS activity and *Y. pseudotuberculosis* virulence [21, 22]. Dynamic changes in pYV PCN and retention of the pYV plasmid during these rapid changes are therefore required for the *Y. pseudotuberculosis* pathogenic lifestyle.

The pYV plasmid encodes several systems important for plasmid regulation and maintenance, including a *cop-rep* locus that regulates plasmid replication, a ParAB partitioning system that ensures each daughter cell inherits the plasmid, and a ParDE toxin-antitoxin (TA) system that promotes plasmid maintenance in the bacterial population [1, 23]. Together, these systems are predicted to synergistically enforce plasmid stability, or the percentage of cells in the population that maintain the plasmid. The highly conserved *cop-rep* locus places pYV in the IncFII group of plasmids [1, 24, 25]. The *cop-rep* locus encodes the essential *<u>rep</u>lication initiation factor* RepA and two negative regulators of RepA expression: the CopA small RNA (sRNA) and the CopB protein. The functions of RepA, CopA, and CopB have been elucidated using plasmid R1, an IncFII plasmid originally isolated from a clinical strain of *S. enterica* that has been extensively studied in *Escherichia coli* model systems [25–27]. These studies support the notion that CopA and CopB regulate the rate of plasmid replication to minimize fluctuations above or below the average PCN set-point [25, 28].

Using *Y. pseudotuberculosis* as a model, we identified the chromosomal *pcnB* gene as a previously unknown regulator of pYV replication and stability. The *pcnB* gene was named for <u>p</u>lasmid <u>c</u>opy <u>n</u>umber <u>B</u> since loss of *pcnB* was found to decrease PCN for several model plasmids in *E. coli* [29, 30]. The *pcnB* gene encodes the poly A polymerase (PAP I), the major bacterial polyadenylase ubiquitous in β- and γ-proteobacteria [29–36]. Polyadenylation, or the addition of adenosine residues to the 3’ OH of an RNA transcript, is poorly understood in bacteria compared to eukaryotes. PAP I-mediated polyadenylation promotes RNA degradation as part of a housekeeping mechanism [37], but is also implicated in certain gene regulatory networks [38–43]. PAP I-dependent polyadenylation has been shown to destabilize RNAs that negatively regulate replication of certain plasmid types in *E. coli*, including CopA from plasmid R1 and RNAI from ColE1 plasmids [29, 44–47]. However, PAP I has not been studied in the context of a naturally occurring virulence or AMR plasmid within its native host, and has never been linked to bacterial virulence.

Here, we report that *Y. pseudotuberculosis pcnB* mutants display reduced PCN and decreased stability of pYV. This defect is associated with reduced T3SS activity in bacterial culture and impaired virulence in a mouse infection model. Furthermore, we show that *pcnB* is required for *Y. pseudotuberculosis* to maintain model AMR plasmids belonging to various plasmid families, highlighting its crucial role in bacterial pathogenicity and antibiotic resistance.

## Results

### *Y. pseudotuberculosis* toggles pYV plasmid copy number in response to temperature

To determine the average PCN per chromosome equivalents at the bacterial population level, we used droplet digital (dd)PCR to measure the number of pYV molecules relative to chromosomes in wildtype *Y. pseudotuberculosis* IP2666pIB1 [22]. Cells were either grown at 26°C to represent growth in environmental conditions outside of the host or at 37°C in low calcium media to represent host cell contact during mammalian infection. The pYV-encoded T3SS is expressed at 37°C and becomes active for secretion following host cell membrane contact or calcium chelation [48–51]. Consistent with previous studies [21, 22], we observed an increase in pYV PCN from an average of ∼1.5 pYV copies per chromosome equivalents at 26°C to ∼3 copies at 37°C/low calcium (Fig 1A). In addition, using a YPIII/pIBX strain of *Y. pseudotuberculosis* harboring a pYV-encoded luciferase reporter, we found that pYV PCN relative to cell density increases one hour after shifting the cultures to 37°C/low calcium conditions (Fig 1B).

**Figure 1.**
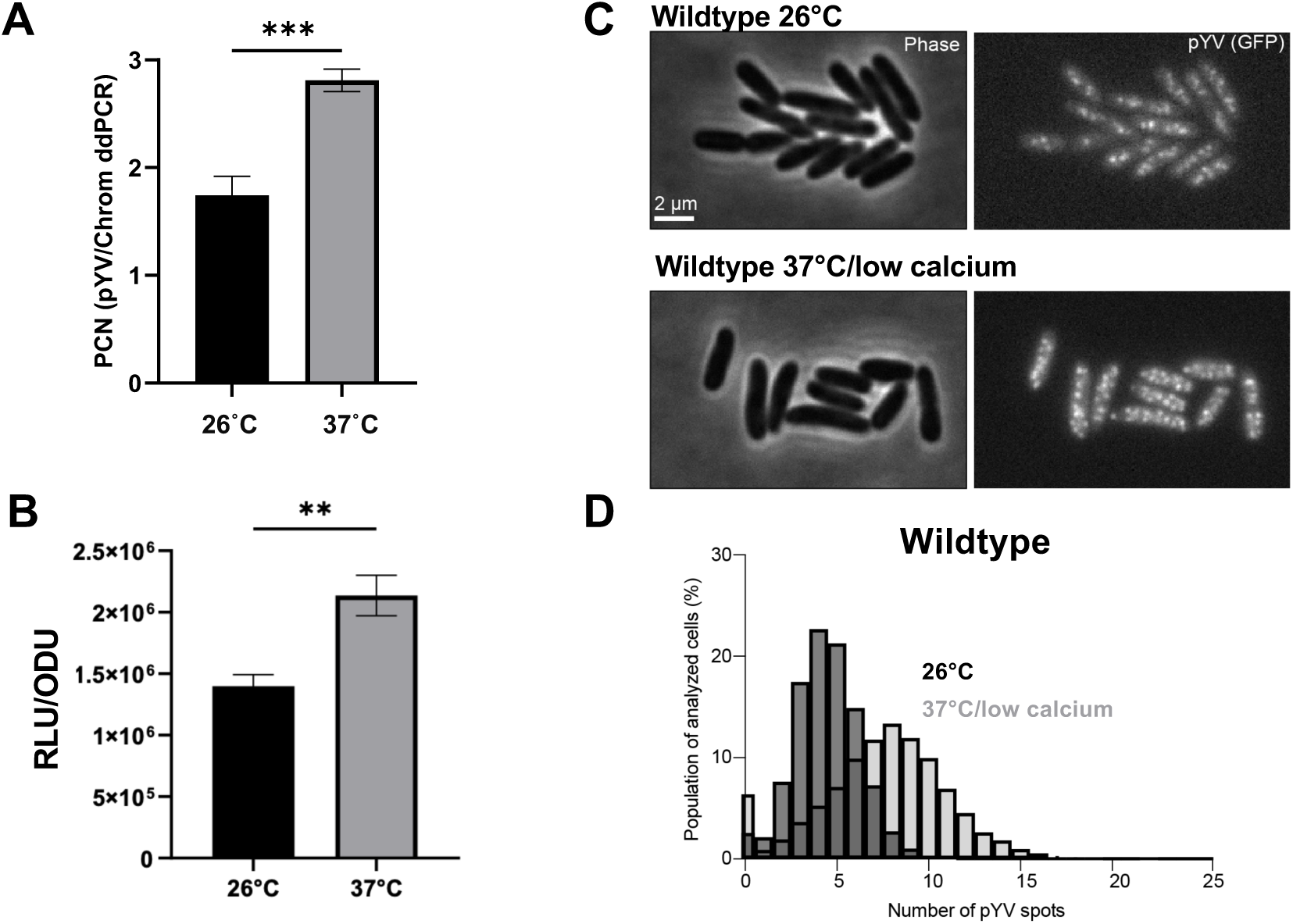
*Y. pseudotuberculosis* pYV plasmid copy number (PCN) dynamically changes in response to temperature. **(A)** pYV plasmid copy number per chromosome was determined for wildtype *Y. pseudotuberculosis* IP2666pIB1 grown at 26°C or 37°C/low calcium using droplet digital PCR (dd)PCR. **(B)** Relative pYV plasmid copy number of wildtype *Y. pseudotuberculosis* YPIII/pIBX was determined following growth at 26°C or 37°C/low calcium using a luciferase reporter stably incorporated in the pIBX plasmid, normalized to bacterial cell density (OD_600_). (A-B) Averages of three independent experiments are shown ± standard deviation (Student t-test, *** p<0.001, ** p<0.01, * p<0.05). **(C)** Representative phase contrast (left) and fluorescence (right) images of *Y. pseudotuberculosis* YPIII/pIBX cells where the pYV plasmid is visualized using a ParB-msfGFP fluorescent marker. Images are shown for cells grown at 26°C or under the 37°C/low calcium condition. **(D)** Histogram depicting the number of ParB-msfGFP-labeled puncta of pYV quantified from data illustrated in panel C. Three biological replicates per condition were imaged and analyzed; for each strain and condition, between 790 to 4428 cells were analyzed per replicate.

To examine pYV PCN at the single-cell level, we employed a microscopy-based approach. To visualize and quantify pYV puncta in living *Y. pseudotuberculosis* cells, we generated a YPIII/pIBX *Y. pseudotuberculosis* strain that harbors a *parB-msfGFP* translational fusion in the native *parB* pYV locus. This approach has been used to visualize plasmids in other bacterial species [52, 53], as ParB binds to plasmid-encoded *parS* sequences but has never been used to study pYV in *Y. pseudotuberculosis*. At 26°C, cells were found to have an average of about four pYV puncta per cell (Fig 1C-D). This number increased to an average of about seven puncta per cell by 1.5 hours after the shift to 37°C (Fig 1C-D). The higher number of pYV per cell than per chromosome equivalents (Figs 1D vs. 1A) is consistent with the expectation that *Yersinia* has more than one chromosome per cell on average under nutrient-rich (fast) growth conditions, similar to *E. coli* [54, 55]. Regardless of the normalization method (per cell or chromosome), *Yersinia* pYV PCN consistently increased by ∼1.5-fold upon shifting bacterial cultures from 26°C to 37°C/low calcium conditions. The pYV plasmid was highly stable in *Y. pseudotuberculosis*, with the vast majority of cells or clones retaining pYV (Fig 1D).

### Disruption of the *pcnB* gene encoding the polyadenylase PAP I decreases pYV PCN and T3SS expression

*Y. pseudotuberculosis* with a functional T3SS, but lacking T3SS Yop effectors (Δyop6), induces activation of the transcription factor NFκB in mammalian cells [56]. The LPS metabolite ADP heptose is delivered to the host cell cytoplasm through the T3SS, stimulating Alpk1/TIFA-dependent NFκB activation [57], although YopJ inhibits NFκB signaling [58, 59]. We used human embryonic fibroblast cells, HEK293T, expressing an NFκB luciferase reporter to identify a *Y. pseudotuberculosis* IP2666pIB1 Δyop6 transposon (Tn) mutant that induced elevated bioluminescence compared to the parental strain, (Δyop6/*pil*::Tn; Fig S1A) [56]. This mutant was found to harbor a Tn insertion in a previously unannotated locus adjacent to the highly conserved pYV *cop-rep* locus. We will refer to this locus as the <u>p</u>lasmid inhibitory locus (*pil*) (Fig S1B). Given the location of the *pil* locus relative to *cop-rep*, we hypothesized that the *pil*::Tn mutation affects the pYV PCN. To test this, we introduced the *pil::Tn* insertion mutation into a clean wildtype *Y. pseudotuberculosis* IP2666pIB1 strain background with intact T3SS effector Yop-encoding genes (referred to as *pil*::Tn). Consistent with our hypothesis, the *pil::Tn* mutant displayed elevated secretion of the T3SS cargo proteins YopE, YopD, and YopH compared to wildtype *Yersinia* (Fig S1C). In addition, we constructed a mutant in the YPIII/pIBX background with a plasmid insertion into the same locus in *pil* (*pil*::pNQ). As expected, both the *pil*::Tn and *pil*::pNQ strains had elevated pYV PCN at 26°C and early after shifting to 37°C relative to the parental strain (Fig S1DE), which is likely the underlying cause of the increased T3SS activity .

Insertion into the *pil* locus was associated with poor growth compared to the parental strain under T3SS-inducing conditions, and suppressor mutants could be readily isolated (Fig S1F). We hypothesized that constitutively elevated pYV PCN contributed to this growth defect, resulting in selective pressure favoring suppressor mutations that would alleviate high pYV PCN. Therefore, we used the *pil* insertion mutation as a tool to identify genes important for pYV PCN homeostasis. We carried out whole genome sequencing of a subset of *pil*::pNQ suppressor mutants and identified several categories of mutations affecting growth, pYV PCN, and the T3SS (Table S1). Two clones harbored a single point mutation that was associated with reduced pYV PCN and T3SS activity relative to the *pil*::pNQ parental strain. This mutation was mapped to the *pcnB* gene encoding the polyadenylase PAP I, and was predicted to confer a leucine to arginine change at position 291 (L291R) in the PAP I neck domain linked to RNA binding (Fig 2A). The L291 residue is conserved in PAP I from *E. coli*, suggesting that the residue might be important for PAP I structure and/or function (Fig 2A). We will use the term *pcnB* when referring to the gene, and PAP I when referring to the *pcnB*-encoded protein.

**Figure 2.**
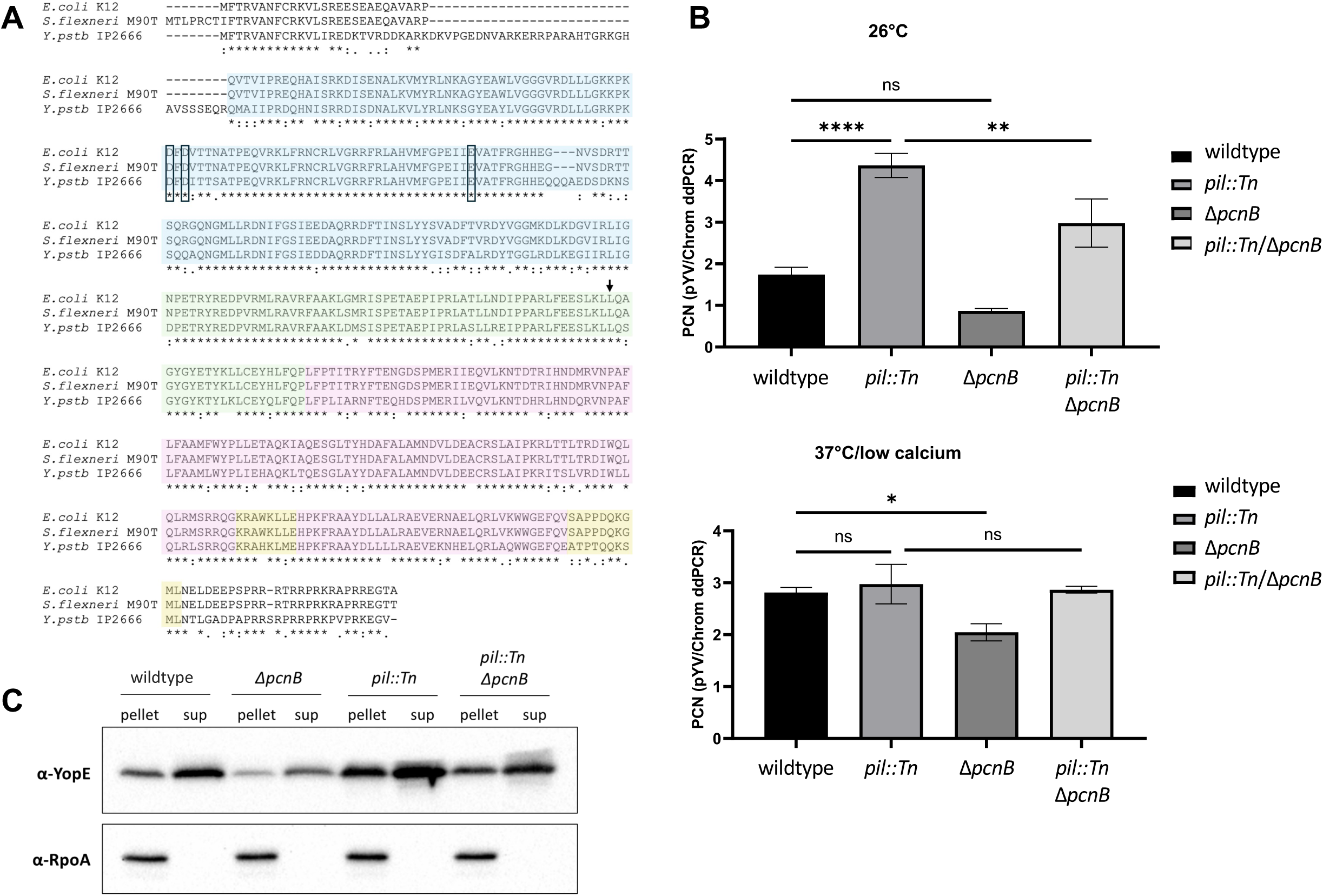
Disruption of the *pcnB* gene encoding poly(A) polymerase PAP I decreases pYV PCN as well as T3SS expression and activity. **(A)** PAP I from *Y. pseudotuberculosis* IP2666pIB1, *E. coli* K12, and *S. flexneri* M90T are highly conserved. Known PAP I domains include the head domain (blue), the neck domain (green), the body domain (purple), and the leg domain (yellow) [62]. Residues in boxes represent known catalytic residues from EcPAP and the residue indicated with an arrow represents the residue that was mutated in the L291R *pcnB* suppressor mutants. **(B)** pYV plasmid copy number per chromosome was determined for *Y. pseudotuberculosis* IP2666pIB1 strains grown at 26°C or 37°C/low calcium using ddPCR. Averages of three independent experiments are shown ± standard error of the mean. Statistical significance was calculated using a one-way ANOVA with Tukey’s multiple comparisons test (**** p<0.0001, *** p<0.001, ** p<0.01, * p<0.05). (C) *Y. pseudotuberculosis* IP2666pIB1 strains were grown in low calcium media at 37°C before preparing samples for western blotting. Bacterial pellets were collected and secreted proteins (Yops) were precipitated from the supernatant (sup). Images shown represent a single blot that was independently probed for YopE and RpoA, which serves as a loading control.

To confirm that the *pcnB* gene encoding PAP I affects *Y. pseudotuberculosis* pYV PCN, we generated a full in-frame deletion of the *pcnB* gene (Δ*pcnB*) in the wildtype and *pil*::Tn IP2666pIB1 strain backgrounds. We used ddPCR to measure relative pYV PCN and found that compared to wildtype, the Δ*pcnB* mutant had lower pYV PCN following growth at 26°C and 37°C/low calcium (Fig 2B). In keeping with our previous findings, the *pil*::Tn mutant displayed significantly higher pYV PCN at 26°C compared to wildtype. Furthermore, loss of PAP I in the *pil*::Tn*/*Δ*pcnB* double mutant significantly reduced pYV PCN at 26°C compared to the *pil*::Tn parental strain. These results show that loss of PAP I leads to decreased pYV PCN. Despite this, loss of *pcnB* did not prevent *Yersinia* from increasing pYV PCN upon shift from 26°C to 37°C, although not to the same high PCN as in wildtype (Fig 2B). Consistent with this reduction in T3SS gene dosage, the Δ*pcnB* mutant displayed two-fold less expression and secretion of the T3SS effector protein YopE compared to wildtype *Y. pseudotuberculosis* (Fig 2C). Similarly, *pcnB* deletion reduced YopE expression and secretion in the *pil*::Tn strain background (Fig 2C). Taken together, these data suggest that *pcnB* is essential for maintaining pYV PCN, and that this function is important for enabling robust T3SS expression under T3SS-inducing conditions.

To determine how the *pcnB* gene and the *pil*::Tn disruption broadly impact expression of pYV-encoded genes, we carried out RNA-seq analyses on wildtype, Δ*pcnB*, *pil*::Tn, and *pil*::Tn*/*Δ*pcnB Y. pseudotuberculosis* IP2666pIB1 grown at 26°C and 37°C/low calcium (Supplementary Dataset 1). Most pYV-encoded gene transcripts were present at higher steady-state level in the *pil*::Tn mutant compared to wildtype *Y. pseudotuberculosis* at both temperatures (Fig S2), consistent with an increase in pYV gene dosage. However, T3SS genes are not translated at 26°C due to the action of several RNA thermometers in *Yersinia* [48], and the *pil*::Tn strain does not display T3SS activity at 26°C despite elevated T3SS gene transcripts at this temperature (Fig S3). Interestingly, the *pil*::Tn strain displayed altered the RNA-Seq read pattern of the region containing *pil*, *copB*, and *repA* at both temperatures (Fig S4). We observed no difference in pYV PCN or T3SS expression when we generated a full in-frame deletion of the predicted ORF found in the *pil* locus (Fig S1C,E). Therefore, we propose that Tn or pNQ insertion into *pil* leads to derepressed transcription of the *cop*-*rep* locus leading to elevated pYV replication through an as-yet unknown mechanism. In contrast, deletion of *pcnB* resulted in fewer transcripts of most pYV-encoded genes compared to wildtype *Yersinia* at both temperatures (Fig S2), consistent with a decrease in pYV gene dosage. In addition, *pcnB* deletion in the *pil*::Tn background also led to a decrease in the steady-state levels of most T3SS genes compared to the *pil*::Tn parental strain (Fig S2). We conclude that loss of *pcnB* leads to a decrease in pYV PCN in both the wildtype and *pil*::Tn backgrounds, leading to decreased transcription of pYV-encoded genes and mitigation of the T3SS activity-induced growth arrest.

### PAP I is required for pYV stability

When working with the Δ*pcnB* strain, we noted that cells lost the pYV plasmid at a higher rate than observed with wildtype *Y. pseudotuberculosis* (Table 1). To determine whether PAP I affects pYV stability in addition to pYV PCN, we generated a Δ*pcnB* strain in the ParB-msfGFP YPIII/pIBX *Y. pseudotuberculosis* background and observed cells lacking clear fluorescent foci under both non-T3SS-inducing (26°C) and T3SS-inducing conditions (37°C, low calcium media. We observed Δ*pcnB* cells with lower cellular GFP fluorescence, consistent with plasmid loss given that the msfGFP-labeled ParB is encoded by pYV (Fig 3A). The presence of these plasmid-devoid cells in the Δ*pcnB* strain resulted in a clear bimodal distribution of cellular fluorescence (Fig 3B), showing that ∼34% of Δ*pcnB* cells lacked pYV (low cellular fluorescence) compared to only ∼3% for the wild-type population at the time of imaging at 26°C (p<10^-100^, Kolmogorov-Smirnov (KS) test; n = 7033 cells per strain). The Δ*pcnB* mutant also displayed a high percentage (24%) of cells lacking pYV under the 37°C/low calcium condition based on the fraction of cell population with low cellular fluorescence (Fig 3B). By analyzing only those cells that retained at least one copy of pYV, we found that the Δ*pcnB* mutant showed the expected increase in ParB-msfGFP puncta per cell from 26°C to 37°C/low calcium (Fig 3B-C). However, the distributions were significantly shifted toward lower values relative to the wildtype strain (p<10^-100^; Fig 3D-E). While the wildtype strain had, on average, 4.5 and 7.9 pYV spots per cell at 26°C and 37°C/low calcium, respectively, the Δ*pcnB* strain had 3.9 and 6.3. Thus, the *pcnB* gene does not affect the ability of the cells to respond to the T3SS-inducing environmental shift, but its loss leads to lower average PCN and increased frequency of plasmidless *Yersinia* cells.

**Table 1.** PAP I is required for pYV maintenance in *Y. pseudotuberculosis*

| Strain: | Percent Retention<sup>□</sup> of pYV after growth at 26°C only: | Percent Retention<sup>□</sup> of pYV after 16 hr exposure to 37°C: |
| --- | --- | --- |
| Wildtype | 100.0 ±0 | 99.1 ±1.2 |
| $\Delta pcnB$ | 81.6 ±17.3 | 57.7 ±10.7 |
| *pil::Tn* | 100.0 ±0 | 99.1 ±1.3 |
| *pil::Tn* $\Delta pcnB$ | 98.1 ±1.8 | 75.9 ±13.1 |

<sup>□</sup> Percent retention of the pYV plasmid was averaged from three independent experiments where ± represents the standard deviation between experiments.

**Figure 3.**
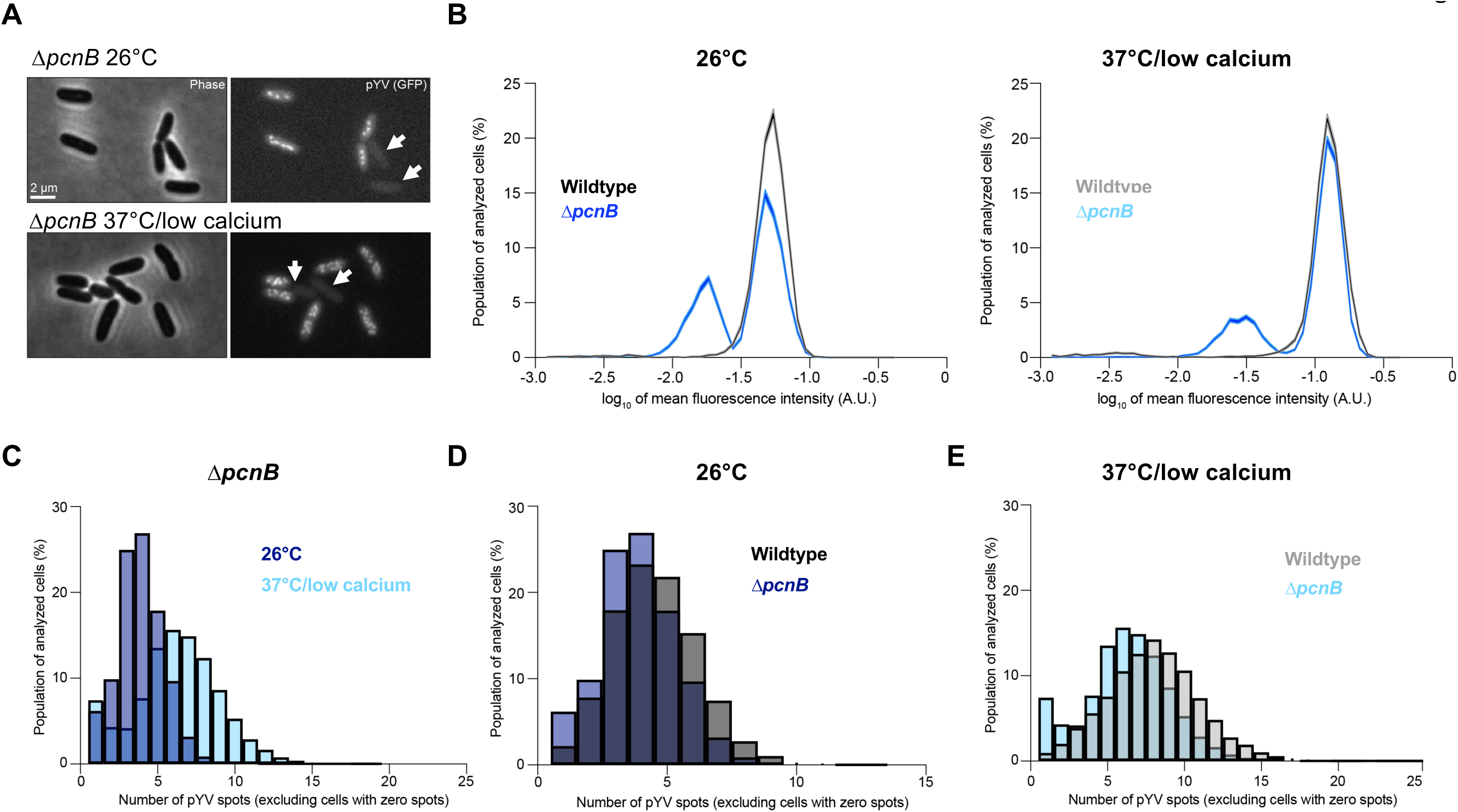
The *pcnB* gene is required for maintenance of the pYV plasmid. **(A)** Representative phase contrast (left) and fluorescence (right) images of *Y. pseudotuberculosis* YPIII/pIBX Δ*pcnB* cells where the pYV plasmid is visualized using a ParB-msfGFP fluorescent marker. Images are shown for cells grown at 26°C or under the 37°C/low calcium condition. White arrowheads indicate plasmidless cells. **(B)** Histograms comparing wildtype vs. Δ*pcnB* cells under 26°C or 37°C/low calcium conditions. Mean fluorescence intensity values are plotted on the x-axis (log_10_). Shaded error bars represent the mean ± standard error of the mean from bootstrapping the data 100 times. Both histograms were randomly sampled without replacement to match the lowest n value of either condition (7,003 and 10,033 for the 26°C and 37°C/low calcium conditions, respectively). Both distributions are significantly different (p < 10^-100^) by Kolmogorov-Smirnov tests. A.U. indicates arbitrary units. **(C)** Histograms depicting the number of ParB-msfGFP-labeled puncta of pYV quantified from data illustrated in panel A. Only cells with at least one detected pYV spot were considered. Three biological replicates were imaged and analyzed per condition. For each strain and condition, between 2063 to 14,479 cells were analyzed per replicate. **(D)** Same as (C) except that the comparison is between wildtype and Δ*pcnB* cells grown at 26°C. For each strain and condition, between 760 to 4179 cells were analyzed per replicate. **(E)** Same as (D) except for the 37°C/low calcium condition. For each strain and condition, between 2046 to 14,479 cells were analyzed per replicate. Distributions in panels D and E are significantly different (p < 10^-80^) by Kolmogorov-Smirnov tests.

To further examine the extent of the pYV plasmid stability defect in the Δ*pcnB* mutant, we passaged bacteria on solid media for several days at 26°C and then assessed pYV retention for each clone by patching onto low calcium media containing the dye Congo red, and incubating at 37°C. Congo red only binds to bacteria that are expressing a functional T3SS [60]. Wildtype cells retained the pYV plasmid at 100% while the Δ*pcnB* mutant cells only retained pYV in 82% of cells following incubation at 26°C for 16 hours during passaging (Table 1). When cells were exposed to T3SS-inducing conditions at 37°C/low calcium and returned to 26°C to mimic the *Y. pseudotuberculosis* facultative lifecycle, we found that only 58% of Δ*pcnB* cells retained pYV compared to 99% for wildtype (Table 1). This observed increase in pYV loss likely stems from the growth arrest of pYV^+^ clones specifically under T3SS-inducing conditions at 37°C [15, 49, 50], which provides a selective growth advantage for Δ*pcnB* clones that have lost pYV. Altogether, these data support a role for PAP I in regulation of pYV PCN and plasmid stability during the *Y. pseudotuberculosis* facultative lifecycle.

### The PAP I^L291R^ mutation affects PAP I protein levels

The original *pcnB* suppressor mutation identified in our *pil*::Tn suppressor screen was a single point mutation that changed leucine in position 291 to an arginine (L291R) (Fig 2A). Although the corresponding L291 residue is conserved in *E. coli*, it has not yet been identified as a critical PAP I residue. Thus, as a first step in investigating the role of this residue, we assessed whether the L291R mutation affects PAP I expression. We introduced the L291R missense mutation into the native *pcnB* locus of both *Y. pseudotuberculosis* IP2666pIB1 (with or without a PAP I C-terminal FLAG tag) and YPIII/pIBX to generate PAP I^L291R^ strains. We found that the PAP I^L291R^ protein was poorly expressed, particularly at 37°C/low calcium (Fig. 4A). These data suggest that the PAP I^L291R^ allele may be unstable, leading to low PAP I expression at least at 37°C/low calcium. Interestingly, a strain engineered with a more conservative leucine to alanine mutation at position 291, PAP I^L291A^, did not significantly affect PAP I expression levels (Fig 4A). Consistent with these data, the PAP I^L291R^ mutant showed enhanced loss of pYV compared to the wildtype strain or the PAP I^L291A^ mutant, as indicated by the emergence of white, fast-growing colonies following growth under 37°C/low calcium conditions on Congo red agar (Fig 4B). Interestingly, the PAP I^L291R^ strain displayed a greater loss of pYV following long passage at 37°C/low calcium compared to when bacteria were passaged at 26°C (Table 2), reflective of the relative stability of the PAP I^L291R^ protein at 26°C. Lastly, the PAP I^L291R^ strain had significantly lower average pYV PCN compared to the wildtype and PAP I^L291A^ strains at 37°C/low calcium (Fig 4C), and this correlated with lower relative T3SS activity (Fig 4D). Taken together, these data suggest that the PAP I^L291R^ mutation disrupts PAP I protein levels, explaining why we identified this mutation in our suppressor screen.

**Figure 4.**
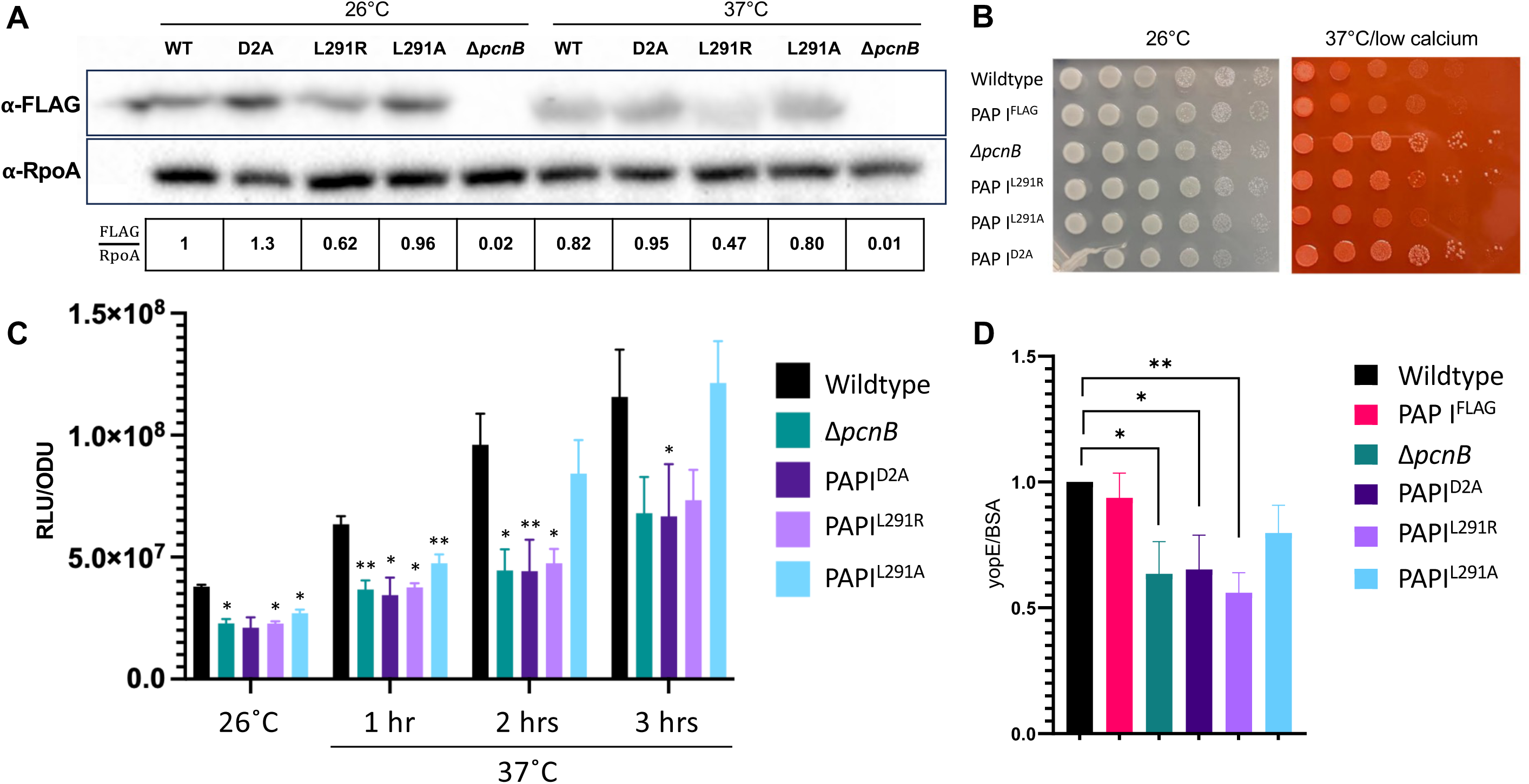
PAP I point mutations lead to a decrease in pYV PCN, plasmid stability, and T3SS activity. **(A)** *Y. pseudotuberculosis* IP2666pIB1 strains harboring various PAP I^FLAG^ alleles grown at 26°C or 37°C/low calcium were pelleted and prepared for western blot analyses. Images shown represent a single blot that was cut after transferring and independently probed for expression of PAP I^FLAG^ (top) and RpoA (loading control, bottom). Data shown is representative of three biological replicates. **(B)** *Y. pseudotuberculosis* IP2666pIB1 strains were grown at 26°C before diluting and spotting onto either LB or low calcium plates containing Congo red dye and incubated at 26°C or 37°C, respectively, prior to imaging. **(C)** Relative pYV plasmid copy number was estimated for strains in the *Y. pseudotuberculosis* pIBX/YPIII background. For each timepoint, luminescence was measured and normalized to cell density (OD_600_). Averages of three independent experiments are shown ± standard error of the mean. Each strain per condition was compared to the wildtype (WT) strain in a one-way ANOVA with Dunnett’s multiple comparisons test (**** P<0.0001, *** P<0.001, ** P<0.01, * P<0.05). **(D)** *Y. pseudotuberculosis* IP2666pIB1 strains were grown at 37°C/low calcium and secreted proteins in the supernatant fractions were precipitated and visualized after separation on as SDS-PAGE gel by Coomassie blue staining. Bovine serum albumin (BSA) was used as a loading and protein precipitation control. The YopE T3SS effector protein band from four independent experiments was visualized, quantified, and normalized to BSA from the same samples. Bars represent the mean fold change in YopE/BSA relative to wildtype and error bars represent the standard error of the mean between different experiments. Statistical significance was determined using a one-way ANOVA with Dunnett’s multiple comparisons test (**** P<0.0001, *** P<0.001, ** P<0.01, * P<0.05).

**Table 2.** Important PAP I residues are required for stability of the pYV plasmid

| Strain: | Percent Retention<sup>□</sup> of pYV after growth at 26°C only: | Percent Retention<sup>□</sup> of pYV after exposure to 37°C: |
| --- | --- | --- |
| Wildtype | 100.0 ±0 | 97.3 ±3.8 |
| $\Delta pcnB$ | 80.2 ±7.8 | 47.7 ±14.0 |
| PAPI<sup>D2A</sup> | 94.0 ±1.7 | 21.0 ±7.0 |
| PAPI<sup>L291R</sup> | 98.6 ±2.5 | 68.6 ±7.9 |
| PAPI<sup>L291A</sup> | 100.0 ±0 | 100 ±0 |

<sup>□</sup> Percent retention of the pYV plasmid was averaged from three independent experiments where ± represents the standard deviation between experiments.

### Mutation of residues predicted to affect polyadenylase activity decreases pYV PCN, plasmid maintenance, and T3SS activity

PAP I D80 and D82 have been shown to be essential for PAP I polyadenylase catalytic activity in *E. coli* [61, 62]. These residues are also present in *Y. pseudotuberculosis* PAP I (Fig. 2A). To determine whether the PAP I polyadenylase activity is responsible for PAP I’s roles in pYV PCN and maintenance, we mutated the two corresponding aspartic acid residues (D114 and D116) to alanine to produce a PAP I^D2A^ catalytic mutant. These mutations did not affect PAP I steady state protein levels (Fig 4A). However, we found that the PAP I^D2A^ mutant displayed reduced pYV PCN at 26°C and 37°C/low calcium compared to wildtype *Y. pseudotuberculosis*, to the same degree as observed in the Δ*pcnB* strain which lacks PAP I completely (Fig 4C). Spot test serial dilution analyses found that PAP I^D2A^ grows similarly to wildtype and Δ*pcnB* at 26°C (Fig 4B). Furthermore, at 37°C/low calcium, PAP I^D2A^ and Δ*pcnB* exhibited about 10-fold more growth compared to wildtype (Fig 4B), consistent with a reduction in growth arrest due to decreased T3SS activity. Similarly, the strain expressing PAP I^D2A^, like the Δ*pcnB* mutant, displayed an increased number of plasmidless, rapidly-growing colonies (Fig. 4B) and reduced pYV stability (Table 2). Lastly, the PAP I^D2A^ allele did not support normal T3SS activity (Fig 4D). Taken together, these data suggest that *Yersinia* PAP I functions as a polyadenylase that maintains pYV PCN, plasmid stability, and T3SS gene expression and activity.

### *Yersinia* PAP I is required for plasmid stability and plasmid-mediated antibiotic resistance from a broad range of plasmid families

To determine if *Yersinia* PAP I contributes to regulation of PCN and retention of other plasmid types, we introduced a ColE1-like pGFP-uv plasmid into wildtype and Δ*pcnB Y. pseudotuberculosis*. This plasmid lacks active partitioning systems such as ParABS, but can be maintained within the population by antibiotic resistance selection [63]. We observed a decrease in fluorescence intensity in Δ*pcnB* colonies compared to wildtype when grown on carbenicillin-containing media, suggesting that lowering of pGFP-uv PCN in the absence of PAP I in *Y. pseudotuberculosis* leads to less robust antibiotic resistance (Fig. 5A). To determine if the decreased PCN of the pGFP-uv plasmid in the Δ*pcnB* strain leads to plasmid loss, we measured kinetics of plasmid retention over 40 hours after removing antibiotic selection. We found that over 70% of Δ*pcnB* cells lose the pGFP-uv plasmid within 10 hours of removing carbenicillin and the plasmid is completely lost from the population by ∼36 hours. In contrast, 100% of wildtype cells retained pGFP-uv over the same 36 hour time course (Fig. 5B).

**Figure 5.**
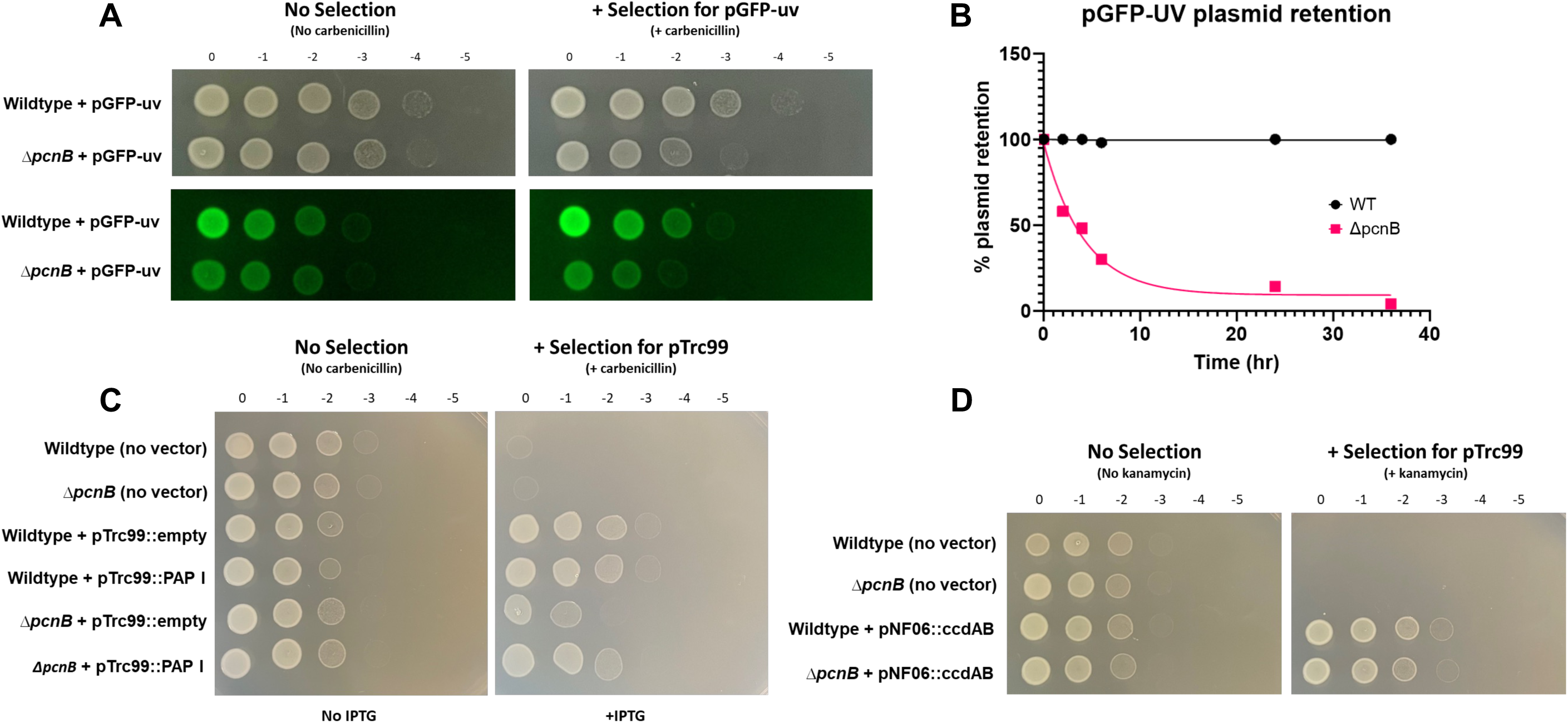
The *pcnB* gene is required for plasmid-mediated antibiotic resistance in *Y. pseudotuberculosis.* **(A)** Wildtype and Δ*pcnB Y. pseudotuberculosis* IP2666pIB1 harboring pGFP-uv were spotted simultaneously onto either plain LB plates or LB plates containing carbenicillin to assess retention of pGFP-uv. Plates were incubated at 26°C for ∼16 hours before imaging spots under a GFP filter. **(B)** Retention of the pGFP-uv plasmid by wildtype and Δ*pcnB Y. pseudotuberculosis* IP2666pIB1 was assessed at 26°C over 40 hours following the removal of carbenicillin. **(C)** Wildtype and Δ*pcnB Y. pseudotuberculosis* IP2666pIB1 harboring either no vector, a pTrc99::empty vector control, or a pTrc99::PAP I recombinant plasmid were spotted onto either plain LB plates or LB plates containing carbenicillin to select for pTrc99 and 0.2 mM IPTG to induce PAP I expression. Plates were incubated at 26°C for ∼16 hours prior to imaging. **(D)** Wildtype and Δ*pcnB Y. pseudotuberculosis* IP2666pIB1 harboring either no vector or a pNF06 mini-F plasmid with a *ccdAB* TA system (pNF06::*ccdAB*) were spotted onto either plain LB plates or LB plates containing 25 µg/mL kanamycin to select for pNF06. Plates were incubated at 26°C for ∼16 hours prior to imaging.

To ensure that the growth defect seen for Δ*pcnB* harboring pGFP-uv was due to loss of PAP I specifically, we next constructed a PAP I over-expression plasmid in the pTrc99 vector backbone. Importantly, pTrc99 is also a ColE1-like plasmid that lacks partitioning systems [64]. We found that Δ*pcnB* harboring pTrc99::empty displays a 10-fold growth defect when spotted onto LB-carbenicillin but not LB-only plates (Fig. 5C), consistent with what was previously observed with the pGFP-uv plasmid. Importantly, this growth defect was nearly completely abolished by expression of PAP I from the pTrc99 plasmid (Fig. 5C). In addition to ColE1 plasmids, we found that PAP I is also required for robust antibiotic resistance conferred by plasmids with p15A-like origins of replication (pACYC184, pBAD33) (Fig S5). Interestingly, replication of plasmids containing ColE-like and p15A-like origins of replication, as well as IncFII plasmids like pYV, is regulated by plasmid-specific sRNA [65–67], suggesting a general role for PAP I in regulating plasmids with replicons regulated by a sRNA. To test if loss of *pcnB* would affect antibiotic resistance conferred by a plasmid whose replication is <u>not</u> known to be regulated by a sRNA, we introduced the mini-F plasmid pNF06::*ccdAB* into wildtype and Δ*pcnB Y. pseudotuberculosis* [30, 44, 68, 69]. The Δ*pcnB* strain grew equally well to the wildtype strain in the presence of antibiotic selection, suggesting that pNF06::*ccdAB* maintenance and copy number is likely not affected by PAP I (Fig. 5D). These data suggest that PAP I is important for *Yersinia* to maintain normal PCN and stability of plasmids whose replication is regulated by a sRNA.

### PAP I is required for *Y. pseudotuberculosis* virulence in mice

It is well established that the pYV plasmid and T3SS are critical for *Y. pseudotuberculosis* virulence in mouse infection models [6, 70]. To determine if loss of PAP I contributes to *Yersinia* virulence, we infected mice via the intraperitoneal (IP) route and monitored colonization of the spleen and liver. Four days post-inoculation, significantly fewer Δ*pcnB*-infected mice exhibited robust colonization of the spleen and liver compared to wildtype-infected mice (Fig 6A and B).

**Figure 6.**
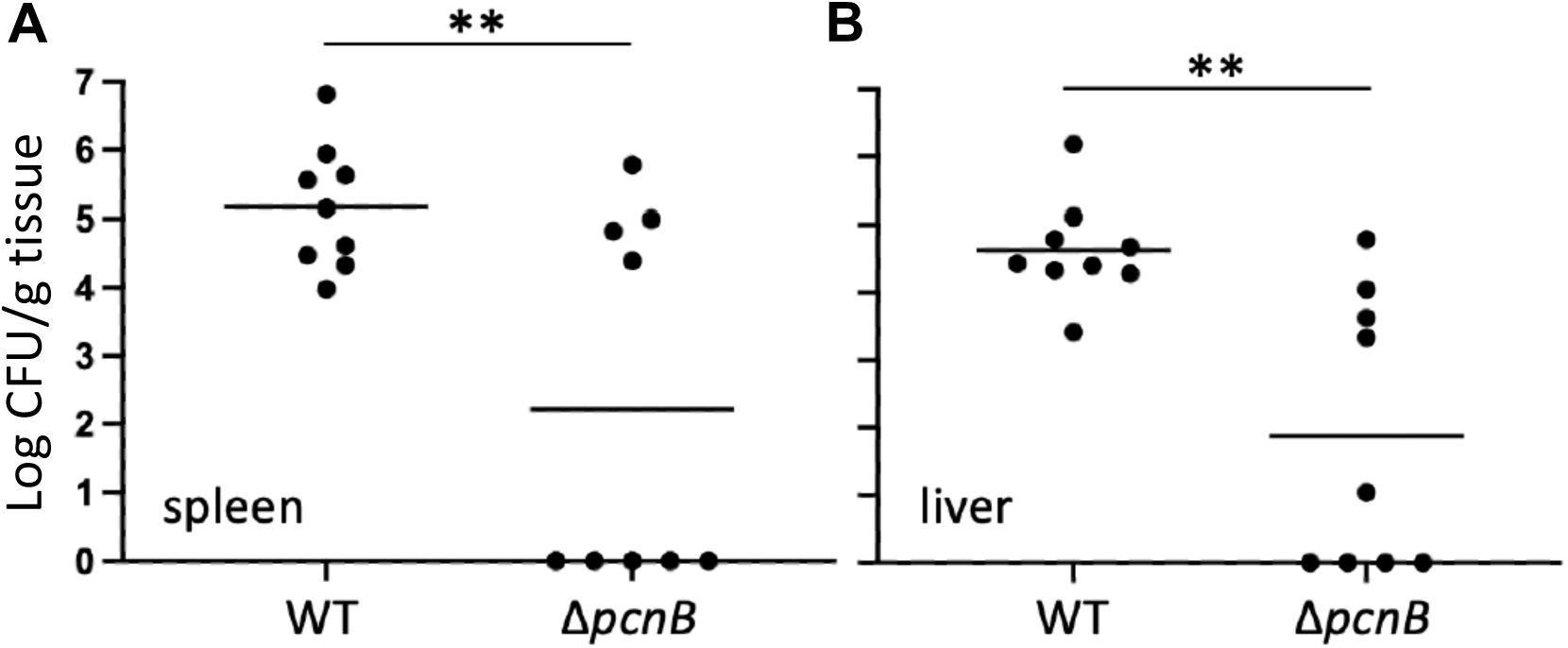
The *pcnB* gene is required for normal *Y. pseudotuberculosis* virulence in a mouse IP infection model. Six-to eight-week-old C57Bl/6 mice were infected with 1-3×10^3^ *Y. pseudotuberculosis* IP2666pIB1 via intraperitoneal (IP) injection. Tissues were collected, homogenized, and plated to determine CFU per gram (CFU/g) tissue 4 days post-inoculation. The log_10_ CFU/g tissue from the spleens **(A)** and livers **(B)** of infected mice are shown. Each circle represents one mouse organ; lines represent geometric means.

Interestingly, Δ*pcnB*-infected mice formed two distinct populations: one population where the spleen and liver were colonized to the same levels as wildtype-infected mice, and a second population that had completely cleared the infection (Fig. 6A and B). We isolated several Δ*pcnB* clones from the livers of colonized mice and found that their pYV stability and T3SS activity phenotypes were similar to the inoculum strain (Fig S6). In addition, we sequenced the genomes of these isolates and could not identify any significant mutations, suggesting that host tissue colonization by the Δ*pcnB* mutant in a subset of mice does not stem from suppressor mutations that stabilize pYV. Taken together, these data suggest that PAP I increases the likelihood of *Yersinia* colonization of host tissues.

### PAP I is required for maintenance of plasmid-encoded T3SS expression and antibiotic resistance in *Shigella flexneri*

The human pathogen *Shigella flexneri* harbors a virulence plasmid, pINV, with a RepFIIA origin similar to the one found on *Yersinia* pYV [1]. To test whether PAP I also regulates *Shigella* pINV, we deleted the *pcnB* gene in the *S. flexneri* M90T Δ*ipaH2.5* (Δ*ipaH2.5::tet^R^*) strain. The resulting Δ*ipaH2.5*Δ*pcnB* mutant was defective in Congo red binding compared to its parental strain (Fig 7A), suggesting a T3SS defect. Indeed, the Δ*ipaH2.5*Δ*pcnB* strain exhibited a stark defect in expression and secretion of the T3SS translocon proteins IpaB, IpaC, and IpaD compared to the parental strain (Fig 7B C). Lastly, we introduced pTRC99 that confers IPTG-inducible expression of *Yersinia* PAP I into the Δ*ipaH2.5* and Δ*ipaH2.5*Δ*pcnB* strains. Consistent with our previous data using *Y. pseudotuberculosis* strains (Fig 5C), deletion of *pcnB* led to a >10-fold loss of antibiotic resistance that could be complemented by PAP I expression (Fig 7D). Taken together, these data suggest that PAP I is essential for maintaining robust plasmid replication, resulting in increased expression of plasmid-encoded virulence and antibiotic resistance genes in pathogenic bacteria.

**Figure 7.**
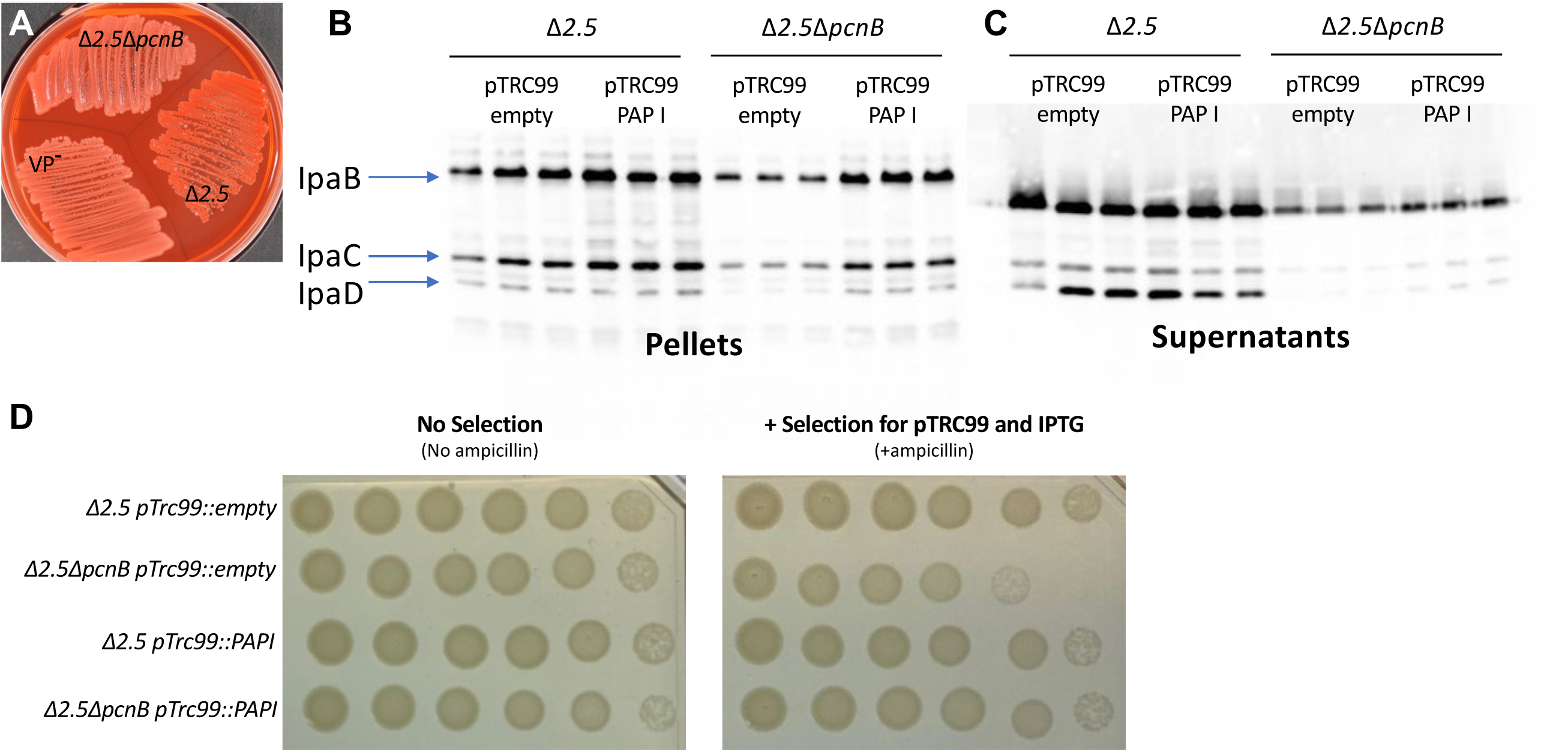
The *pcnB* gene is required for plasmid-encoded expression of T3SS and antibiotic resistance genes in *Shigella flexneri*. **(A)** *S. flexneri* M90T Δ*ipaH2.5* and Δ*ipaH2.5*Δ*pcnB*, as well as a virulence plasmid cured (VP^-^) BS103 strain, were incubated on Congo red media at 37°C. **(B)** *S. flexneri* M90T Δ*ipaH2.5* and Δ*ipaH2.5*Δ*pcnB* carrying pTRC99 with and without IPTG-inducible *Yersinia* PAP I were incubated at 37°C, and secreted T3SS effector proteins were visualized after separation on as SDS-PAGE gel using anti-IpaB, anti-IpaC, and anti-IpaD antibodies. **(C)** *S. flexneri* M90T Δ*ipaH2.5* and Δ*ipaH2.5*Δ*pcnB* carrying pTRC99 with and without IPTG-inducible *Yersinia* PAP I were grown on either plain TCS media or TCS containing ampicillin and IPTG. Shown is one of two independent experiments.

## Discussion

*Y. pseudotuberculosis* modulates the PCN of its pYV virulence plasmid to control the dosage of pYV-encoded genes in response to temperature [21, 22]. This ability of *Y. pseudotuberculosis* to upregulate pYV PCN at mammalian body temperature is critical for its ability to cause disease. Here, we show that in addition to dynamic regulation of pYV PCN, *Y. pseudotuberculosis* requires the chromosomal gene *pcnB*, encoding the PAP I polyadenylase, to maintain pYV PCN homeostasis and prevent pYV loss from the bacterial population. The role of the *pcnB* gene in promoting virulence plasmid-encoded T3SS expression is conserved in both *Y. pseudotuberculosis* and *S. flexneri*. Although *pcnB*/PAP I has been shown to regulate plasmid replication of model plasmids in *E. coli*, this is the first study to link this gene to bacterial pathogenesis. In addition, the requirement of *Yersinia* and *Shigella pcnB*/PAP I for robust plasmid-encoded antibiotic resistance suggests that PAP I represents a potential new drug target.

Putative PAP I orthologs have been bioinformatically identified in many bacterial species hailing from diverse phyla, and yet studies on *pcnB*/PAP I have historically been confined to a small number of non-pathogenic bacterial species [36]. *E. coli* PAP I (EcPAP) has been shown to promote RNA degradation of target transcripts. Degradation of RNA is generally initiated by the endonuclease RNase E and, in the absence of 3’ secondary structures, the RNA fragments produced by RNase E are then digested by the 3’-5’ exonulceases PNPase and RNase II [71]. However, RNase II-and PNPase-mediated RNA degradation is significantly impaired for transcripts that have terminal stem-loop structures, such as mRNAs containing intrinsic terminators at their 3’ ends and structured sRNAs [71]. PAP I has been shown to add poly(A) tails to the 3’ ends of transcripts containing terminal stem-loops structures to promote their decay [72–74] and it is thought that PAP I-dependent polyadenylation of these transcripts provides a “toe-hold” for PNPase and RNase II to bind and digest the RNA [74–77]. Components of the RNA degradosome such as PNPase are known to be important for T3SS function in *Y. pseudotuberculosis* [78–80] and PAP I has been shown to copurify with PNPase in *E. coli* [81], suggesting that PAP I may represent a novel component of the *Yersinia* RNA degradosome.

Importantly, many of the major domains and catalytic residues are conserved between *Yersinia* and *Shigella* PAP I and EcPAP. EcPAP polyadenylase activity is abolished when the two catalytic residues Asp80 and Asp82 are mutated to alanine residues in *E. coli* [61, 62]. To test if the phenotypes of the Δ*pcnB* strain was due to loss of PAP I polyadenylase activity, we mutated the corresponding catalytic residues (Asp114 and Asp116) in PAP I from *Y. pseudotuberculosis* to generate a PAPI^D2A^ strain with both catalytic residues replaced by alanine residues. The PAPI^D2A^ mutant recapitulated the phenotypes observed for the Δ*pcnB* strain, including reduced pYV PCN, plasmid stability, and T3SS activity. This supports that PAP I acts as a polyadenylase to regulate plasmid maintenance and T3SS activity in *Y. pseudotuberculosis*, and our data suggests that this may also be the case in *S. flexneri*.

EcPAP has been shown to polyadenylate both chromosomally- and plasmid-derived transcripts and was first identified due to its regulation of a sRNA that regulates plasmid replication [29, 30]. EcPAP has been shown to regulate PCN for the IncFII family plasmid R1 by polyadenylating and targeting CopA regulatory sRNA for degradation, leading to de-repression of R1 plasmid replication and increasing R1 PCN (46). This is consistent with our finding that PAP I is required for normal PCN and maintenance of the IncFII family *Yersinia* pYV plasmid, and for normal expression of the *Shigella* T3SS encoded on the related pINV virulence plasmid [1]. Taken together, these data point to the possibility that PAP I targets CopA sRNA from the *Yersinia* pYV and *Shigella* pINV *cop-rep* loci for polyadenylation. Bacteria are thought to maintain large extrachromosomal elements such as IncF virulence plasmids within a tight PCN range near a specific “set point” that is optimal for that species lifestyle [25]. This is particularly complicated in the case of pYV since this plasmid seems to have a different optimal PCN “set point” depending on temperature. Given that the Δ*pcnB* mutant displays a PCN defect at both 26°C and 37°C, this suggests that pYV plasmid replication rates are unusually low in the absence of PAP I in general, possibly due to the buildup of CopA sRNA. Reduced plasmid replication rates can also lead to plasmid stability defects when the plasmid replication rate drops low enough that cells can no longer reliably pass a copy of the plasmid to daughter cells during cell division [82]. Both reduced PCN and reduced plasmid stability can theoretically drive compounding disturbances in plasmid maintenance systems due to reduced gene dosage of stability systems such as partitioning and toxin-antitoxin (TA) systems. In keeping with this idea, our RNA-seq analysis supports the notion that the decrease in pYV-encoded gene dosage in the absence of PAP I leads to decreased expression of the ParDE TA system in Δ*pcnB* cells that may be responsible for killing plasmidless cells (Supplementary Dataset 1)[23]. Altogether, this supports the claim that PAP I is required for maintenance of the pYV PCN “set point” at both 26°C and 37°C, possibly through modulation of CopA sRNA levels. This, in turn, may decrease the frequency at which plasmidless daughter cell arise. Alternatively, PAP I may also regulate stability of additional transcripts that control plasmid partitioning and/or post-segregational killing. Future studies will focus on identification of polyadenylated transcripts in *Yersinia* and *Shigella*.

While there is an overall reduction in pYV PCN in the absence of *pcnB*/PAP I, Δ*pcnB* cells that retain the pYV plasmid still increase pYV PCN upon shifting from 26°C to 37°C. In addition, we have found that the *pcnB* gene is transcribed at 26°C and 37°C (Supplementary Dataset 1) and the PAP I protein is also produced at both temperatures (Fig 4A). Taken together, this suggests that PAP I is not required for the temperature-dependent shift in pYV PCN but rather for the maintenance of the pYV plasmid in general as it undergoes dynamic shifts in PCN during the *Yersinia* facultative lifestyle. We hypothesize that PAP I polyadenylates and promotes degradation of pYV-encoded CopA sRNA to regulate cellular CopA levels at both 26°C and 37°C. This would help maintain CopA levels within a range that promotes maintenance of pYV PCN near its optimal “set point” and ensuring that pYV-encoded gene dosage remains high enough to support replication and maintenance of the plasmid under conditions where pYV PCN is normally kept low. In the absence of PAP I, we hypothesize that pYV plasmid replication rates decrease due to abnormally high CopA sRNA levels, leading to reduced pYV PCN, plasmid stability, T3SS activity, and host tissue colonization.

We found that PAP I is required for normal *Y. pseudotuberculosis* virulence in a mouse intraperitoneal infection model, consistent with its role in promoting high pYV-encoded gene dosage and pYV stability, and therefore T3SS activity. Interestingly, Δ*pcnB*-infected mice fell into two groups: one in which the spleen and liver were colonized with *Yersinia* to wildtype-infected levels, and another in which no Δ*pcnB* cells could be recovered from spleens and livers four days post-inoculation. Overall, this led to a significant decrease in spleen and liver colonization by Δ*pcnB* compared to wildtype *Y. pseudotuberculosis*. Consistent with our finding that deletion of *pcnB* leads to enhanced pYV loss from the bacterial population *in vitro*, wildtype cultures used for mouse inoculation had 100% retention of pYV whereas the Δ*pcnB* inoculum had an average retention of 81% at the time of inoculation. This 19% decrease in pYV+ Δ*pcnB* cells in the inoculum are not likely to explain the difference in virulence, as wildtype inoculums of 10^3^ to 3×10^3^ yielded similar spleen and liver colonization levels. These data suggest that *pcnB*/PAP I may aid *Yersinia* surviving a colonization bottleneck during infection. To our knowledge, this is the first time that PAP I has been directly linked to virulence of a bacterial pathogen and highlights the importance of understanding how PAP I and virulence plasmid gene dosage contribute to bacterial pathogenesis. All Δ*pcnB* cells recovered from the spleen and livers of infected mice retained at least one copy of the pYV plasmid despite the plasmid stability defect observed for this strain *in vitro*. These data suggest a selective pressure to retain pYV during growth in mouse tissues, consistent with the importance of the T3SS to *Yersinia* virulence. This contrasts with what we observe *in vitro* when pYV and T3SS expression is a metabolic burden and not a benefit. These data suggest that *pcnB*/PAP I is required for maintenance of the pYV plasmid when pYV is dispensable to survival. This is necessary to ensure that enough of *the Y. pseudotuberculosis* population is pYV^+^ to be able to colonize a new host and survive in host tissues.

In addition to pYV and pINV, we found that *pcnB* is important for robust replication of sRNA-regulated plasmids encoding antimicrobial resistance (AMR) genes in *Yersinia* and *Shigella*. When *Y. pseudotuberculosis* or *S. flexneri* Δ*pcnB* cells are grown in the presence of the plasmid-selective antibiotic, we observed a ten-fold decrease in growth (Fig 5 and 7), consistent with previous studies in *E. coli* [29]. Similarly, we found a ten-fold decrease in GFP fluorescence in the *Y. pseudotuberculosis* Δ*pcnB* strain carrying the ColE1 pGFPuv plasmid compared to wildtype, suggesting a ten-fold decrease in plasmid-encoded gene expression in the absence of *pcnB*. In addition, time course experiments found that the majority of the Δ*pcnB* population loses the pGFPuv plasmid within just 10 hours of removing the selective antibiotic. These data support that ColE1 PCN and plasmid stability are reduced in the absence of PAP I, rendering cells more antibiotic susceptible. Interestingly, the rate at which pYV is lost from *Y. pseudotuberculosis in vitro* under non-T3SS inducing conditions was far slower than for pGFPuv. We passaged wildtype and Δ*pcnB* strains in LB medium at 26°C and while 100% of wildtype retained pYV, only ∼60% of Δ*pcnB* cells retained pYV by 13 days with a still increasing rate of loss. As pGFPuv has no active plasmid partitioning or maintenance systems but pYV has both ParABS and ParDE systems, we hypothesize that the rate of plasmid loss in cells lacking PAP I activity will depend on the presence and type of plasmid-encoded mechanisms that promote plasmid stability.

Through our RNA-seq analysis we identified several differentially regulated transcripts in the *Y. pseudotuberculosis* Δ*pcnB* mutant that are involved in responding to several types of stress, including the universal stress protein UspB, the *Yersinia* urease operon, and several cold-shock proteins (Supplemental Dataset 1). Importantly, these differentially regulated genes are encoded on the *Y. pseudotuberculosis* chromosome and so should not be significantly affected by the pYV PCN defect in the Δ*pcnB* mutant. In addition, EcPAP from *E. coli* has been previously reported to play a critical role in regulating mRNA transcripts important for responding to stress where loss of PAP I leads to stabilization of stress response transcripts [43]. Deletion of *pcnB* in *E. coli* results in increased tolerance to several stresses, including osmotic stress, acid stress, heat shock, and cold shock [83]. Interestingly, we found that loss of PAP I does not affect *Y. pseudotuberculosis* viability following heat shock, growth at 4°C, acid shock, or osmotic shock (Fig S7). Our RNA-seq analysis also found that some stress response genes such as CspE are downregulated in Δ*pcnB* cells, suggesting that PAP I normally promotes stability of at least some stress-related transcripts in *Y. pseudotuberculosis*. These data suggest that while PAP I stabilizes sRNA-regulated plasmids in multiple bacterial species, its effect on stress responses is species-specific.

While *E. coli* EcPAP and PAP I from *Y. pseudotuberculosis* and *S. flexneri* are highly conserved in general, some regions of the proteins are less conserved and might account for their different impacts on stress tolerance. Interestingly, while the PAP I N-terminal region following the initial propeptide sequence is predicted to be disordered, this is also the region that displays the least amount of conservation between the three proteins. In contrast, this N-terminal region is 100% conserved between *Y. pseudotuberculosis* and *Y. pestis* (Fig S8). This disordered region is also significantly expanded in length for PAP I from *Y. pseudotuberculosis* and *Y. pestis* compared to *E. coli* and *Shigella* PAP I (Fig 2A). Given that disordered regions of RNA-binding proteins have been proposed to mediate interactions with target RNAs and/or other proteins [84], it is possible that *Yersinia* PAP I has evolved to target different RNA transcripts and/or interacts with one or more unique protein partners compared to EcPAP. *Yersinia* PAP I complements the plasmid-encoded antibiotic resistance defect of the *Shigella* Δ*pcnB* mutant, suggesting that PAP I plays a conserved role in regulating plasmid-derived sRNAs. However, given the different role of PAP I in stress resistance in *Yersinia* and *E. coli*, PAP I may be adapted to regulate chromosomally-derived mRNA transcripts in a more species-specific way.

In this study, we show for the first time that *pcnB*/PAP I is required for maintenance of a large IncFII virulence plasmid (pYV) that undergoes dynamic yet specific changes in PCN to regulate virulence gene dosage in *Y. pseudotuberculosis*. PAP I was also found to be required for normal *Y. pseudotuberculosis* virulence, providing the first direct evidence that PAP I regulates virulence of a Gram-negative pathogen. PAP I is required for maintenance of plasmids from multiple families in *Y. pseudotuberculosis* and *S. flexneri* (Fig 5 and 7; Fig S5), including families that commonly include virulence and antimicrobial resistance plasmids. PAP I homologs can be found in many species of Gram-negative pathogenic bacteria, many of which harbor extrachromosomal plasmids important for their pathogenic lifestyles, and genomic studies show that the *pcnB* gene is highly subject to horizontal gene transfer [36]. Our data support the idea that bacterial PAP I has a conserved role in promoting the stability of many sRNA-regulated plasmids and may be critical for the establishment of new plasmids in a bacterial population following horizontal transfer by promoting plasmid replication. However, given the different effects of PAP I loss to stress resistance in *E. coli* and *Yersinia*, PAP I-mediated polyadenylation of transcripts that are chromosomally-encoded likely varies across species. Future studies will explore the possibility of targeting PAP I activity as a novel antibiotic and/or plasmid-curing strategy as well as the contribution of PAPI-mediated polyadenylation to regulation of gene expression in different pathogenic bacteria.

## Materials and Methods

### Bacterial strains, plasmids, and growth conditions

All strains used in this study are listed in Table S2. *Y. pseudotuberculosis* strains were grown in LB or 2xYT media (pH=7) at 26°C or 37°C shaking at 250rpm, unless otherwise indicated. *S. flexneri* were grown in tryptic soy broth media at 30°C or 37°C on a roller.

### Construction of *Y. pseudotuberculosis* and *S. flexneri* mutant strains

All primers and plasmids used to generate *Y. pseudotuberculosis* strains for this study are listed in Tables S3 and S4, respectively. Inserts were amplified by PCR using primers designed for Gibson Assembly and final recombinant plasmids were all made using the pCVD442 suicide vector backbone (λpir-dependent replicon, carbenicillin resistant [Carb^R^], *sacB* gene conferring sucrose sensitivity) (Addgene) following digestion with NdeI and BbsI. All ligations were performed using NEBuilder HiFi DNA Assembly kit (New England Biolabs). Recombinant pCVD442 plasmids were transformed into *E. coli* S17-1 λpir competent cells and introduced into Y*. pseudotuberculosis* IP2666pIB1 or YPIII/pIBX via conjugation. Carb^R^ and irgansan-resistant (*Yersinia* selective antibiotic) clones were grown in the absence of antibiotics and plated onto media containing 10% sucrose to induce SacB-mediate toxicity for pCVD442 loss. Carb sensitive, sucrose-resistant, and Congo red-positive (pYV+) clones were screened by PCR and sequence verified by Sangar sequencing. *<u>pil</u>* <u>insertion strains:</u> A *pil::Tn Y. pseudotuberculosis* IP2666pIB1 Δyop6 strain was originally isolated from a Tn*Himar1* library screened for altered NFκB induction compared to the parental strain (52). The Tn insertion was then transferred to the wildtype IP2666pIB1 strain using the pSR47s suicide plasmid, as described above. The *pil::pNQ Y. pseudotuberculosis* YPIII/pIBX mutant was generated using the synthesized pNQ705 suicide plasmid carrying a truncated fragment of *pil* (Genescript)[85]. <u>Δ*pcnB*:</u> ∼500 bp upstream of *pcnB* and ∼500 bp downstream of *pcnB* was PCR amplified using genomic DNA (gDNA) from *Y. pseudotuberculosis* IP2666pIB1. Amplified fragments were cloned into pCVD442 to produce pCVD442::Δ*pcnB*. <u>ParB-GFP:</u> To make ParB-GFP YPIII/pIBX strains, three fragments were amplified for insertion into pCVD442: (1) the last ∼500 bp of the *parB* gene excluding the stop codon was amplified from *Y. pseudotuberculosis* gDNA, (2) msfGFP was amplified from pDK13 and (3) 500 bp downstream of the *parB* gene was amplified from *Y. pseudotuberculosis* gDNA. The resulting fragments were cloned into pCVD442 to generate a *parB*-msfGFP construct that includes 500 bp of downstream homology after msfGFP. <u>PAPI-C-6xHis and PAPI-C-3xFLAG:</u> To make PAPI-6xHis strains, *pcnB* was amplified from *Y. pseudotuberculosis* gDNA and cloned into pET28 that was previously digested with Xhol and NcoI to produce PAPI-C-6xHis. PAPI-C-6xHis was then amplified from pET28 and 500bp downstream of *pcnB* was amplified from gDNA for insertion into pCVD442 to make pCVD442::PAPI-c-6xHis containing 500 bp of homology flanking the 6xHis tag. We subsequently found that the 6xHis tag was not sufficient to visualize the PAPI protein, and so alternative PAPI-C-3xFLAG strains were made in order to carry out western blot analyses. To make PAPI-3xFLAG, three fragments were amplified for insertion into pCVD442: (1) the *pcnB* gene was amplified from *Y. pseudotuberculosis* gDNA (2) 3xFLAG was amplified from pET28 and (3) 300 bp downstream of *pcnB* was amplified from *Y. pseudotuberculosis* gDNA. The fragments were then cloned into pCVD442 to make pCVD442::PAPI-c-3xFLAG containing 300 bp of downstream homology following the 3xFLAG tag. <u>PAP 1 point mutations:</u> To make PAP I^D2A^, PAP I^L291R^, and PAP I^L291A^ mutants, we used the Q5 site-directed mutagenesis kit (New England Biolabs) to introduce base pair changes at the corresponding residues using the pCVD442::PAPI-c-6xHis or pCVD442::PAPI-c-3xFLAG recombinant plasmids as templates.

<u>Δ*ipaH2.5*Δ*pcnB Shigella*</u>: A fragment containing 200 bp of homology upstream and 240bp of downstream *pcnB* flanking sequence was amplified from ECK0142 (<u>Δ*pcnB*</u> *E. coli* K-12 BW25113) (PMID 16738554) using pcnb_F and pcnb_R. This fragment was introduced into the chromosome of Δ*ipaH2.5::tetR Shigella* (PMID: 25378474) using lambda red recombination (PMID: 10829079). The insertion was confirmed using pcnB_R_op and k2_wanner.

### *In vitro* suppressor screen

LB cultures of the *pil::*pNQ YPIII/pIBX mutant strain were grown at 26°C shaking overnight before diluting in fresh media and spotting for single colonies onto low calcium plates (LB supplemented with 20 mM sodium oxalate and 20 mM MgCl_2_). Plates were incubated at 37°C for ∼16 hours to induce T3SS expression and selective pressure for growth of suppressor mutants. Clones that broke growth arrest and thus formed large colonies were selected and screened for changes in pYV PCN using the luciferase assay (see below). Suppressor mutants were also screened for changes in T3SS activity using an *in vitro* secretion assay (see below).

### Identification of mutations arising from *pil::*pNQ suppressor screen

A subset of *pil::*pNQ YPIII/pIBX suppressor mutants were selected for whole genome sequencing. In brief, cells were grown shaking at 26°C overnight and total genomic DNA was extracted using the Blood & Tissue DNeasy kit (Qiagen) according to manufacturer recommendations. Qubit quantification, library preparation, and Illumina paired-end sequencing with 150 bp read length was carried out by the UC Davis DNA Technologies Core. Adapters and low-quality reads were trimmed by Trimmomatic (Version 0.39)[86]. The trimmed sequences from each sample were aligned to the reference genome using BWA [87] The reference sequence was composed of the YPIII chromosome (CP009792.1) [88] and its plasmid (CP032567.1)[89]. Samtools [90] was used to sort and index the aligned reads. Duplicate reads originated from a single DNA fragment during sample preparation were located and tagged by MarkDuplicates in Picard (http://broadinstitute.github.io/picard/). Single nucleotide polymorphisms were detected and joined by GATK (Version 4.1.6) [91].

### Droplet digital PCR (ddPCR)

ddPCR to determine pYV plasmid copy number was carried out as previously described [92]. Total DNA was extracted from bacterial cells using GeneJet Genomic DNA Purification kit (Thermo Scientific) according to the manufacturer’s recommendations for Gram-negative bacteria. Each genomic DNA sample was split into two reactions and the primer pair F_chrom_ddPCR/R_chrom_ddPCR was used for to amplify chromosomal DNA and F_pYV_ddPCR/R_pYV_ddPCR was used to amplify pYV DNA. Individual ddPCR samples were prepared as follows: 10.5 µL of total genomic DNA was combined with 12.5 µL of 2xEVA-Green master mix for unlabeled probes (Bio-Rad), 1 µL of HindIII-HF (NEB), and 1 µL of respective primer pairs (200 nM final concentration) in 8-strip PCR tubes. PCR tubes were incubated at room temperature for 10 minutes to allow HindIII to digest genomic DNA. Droplet generation was carried out using an Automated Droplet Generator (Bio-Rad) and droplets were subsequently transferred to a deep-dish qPCR plate (Bio-Rad) for PCR amplification. PCR was carried out with hot-start activation at 95°C for 5 mins following by denaturation at 94°C for 30s, amplification at 58°C for 1 min over 40 cycles, and signal stabilization at 4°C for 10 min and 90°C for 5 min. A ramp rate of 2C/s was used for all PCR steps. Droplets were analyzed using a QX200 droplet reader (Bio-Rad) and data was analyzed using QuantaSoft Analysis Standard Edition (version 1.2) (Bio-Rad). Statistical significance was calculated using a one-way ANOVA with Tukey’s multiple-comparisons test (**** P<0.0001, *** P<0.001, ** P<0.01, * P<0.05). All graphs and statistical analyses were generated using GraphPad Prism (10.2.0).

### Luciferase pYV plasmid copy number assay

Overnight cultures for strains in the YPIII/pIBX background were grown shaking at 26°C before subculturing to an OD_600_ of 0.2 in 2xYT low calcium media (containing 20mM sodium oxalate and 20mM MgCl_2_). Cultures were then grown shaking for 1.5h at 26°C before shifting to 37°C. Luminescence and OD_600_ were measured after 1.5h at 26°C just before shifting to 37°C and every hour after shifting to 37°C. Samples were collected at each timepoint in white walled, clear bottom 96 well plates and luminescence (using lum aperture for 96 well plate with measurement time=0.1s) and OD_600_ (using Photometric 600 excitation filter) was measured using a PerkinElmer Multimode Plate Reader EnVision system. Relative plasmid copy number was determined by dividing luminescence by OD_600_ for the same sample. Statistical analysis was determined using a one-way ANOVA with Dunnett’s multiple comparisons test (**** P<0.0001, *** P<0.001, ** P<0.01, * P<0.05) where applicable. All graphs were made using GraphPad Prism (10.2.0).

### ParB-msfGFP microscopy and image analysis

Wildtype and Δ*pcnB* YPIII/pIBX cells expressing ParB-msfGFP from LB plates supplemented with kanamycin were inoculated into EZ Rich Defined Medium (EZRDM) (Teknova) without antibiotic supplementation. EZRDM medium was chosen to reduce medium autofluorescence for downstream imaging purposes. Cultures were grown shaking at 200 rpm overnight at 26°C to a final OD_600_ of ∼2-4. A secretion analysis was performed to confirm that *Y. pseudotuberculosis* displays normal T3SS activity when grown in EZRDM under T3SS-inducing conditions (37°C, supplemented with 20 mM sodium oxalate and 20 mM MgCl_2_), referred to as low calcium EZRDM (Fig S9). For non-T3SS inducing conditions, overnight cultures were diluted to an OD_600_ of ∼0.2 in fresh EZRDM and grown shaking for 3 hours at 26°C. For T3SS-inducing conditions, overnight cultures were diluted to an OD600 of ∼0.2 in low calcium EZRDM and grown shaking for 1.5 hours at 26°C before shifting to 37°C for an additional 1.5 hours.

Cells were prepared for imaging by spotting onto a 1% agarose-low calcium EZRDM pad and covering with a no. 1.5 coverslip. Imaging was carried out using a Nikon Eclipse Ti microscope equipped with a 100X Plan Apo 1.45 NA phase-contrast oil objective, a Hamamatsu Orca-Flash4.0 V2 CMOS camera, and CoolLED pE-400 (Lumencor). The microscopes were controlled by the Nikon Elements software. The following Chroma filter cubes were used to acquire the fluorescence images: GFP: excitation ET470/40x, dichroic T495lpxr, emission ET525/50Lm. Images were acquired with 50 ms of exposure (50% power) for GFP and 100 ms for phase contrast. For each strain and condition, three independent biological replicates (inoculated with different colonies) were grown and imaged.

For image analysis, cells were segmented and cell outlines were generated using the open-source Oufti software [93] using the following parameters: Edgemode, 1; Dilate, 1; openNum, 0; InvertImage, 0; ThreshFactorM, 0.86254; ThreshMinLevel, 0.3; EdgeSigmaL, 0.5; LogThresh, 0. Cell meshes were further curated using the SVMCuratedCellDetection.m script to identify and remove misdetected cells [94]. To quantify the number of pYV copies per cell, detection of pYV spots was performed using the ImageJ plug-in package ThunderSTORM [95, 96]. For of cells grown at 26°C, image filtering used a wavelet filter (B-spline) with a scale of 2.0 and order of 3.0. Under non-T3SS-inducing conditions, a local maximum using 5.0*std(Wave.F1) and a connectivity of 8-neighborhood was used to localize molecules based on peak intensity thresholds, and sub-pixel localization used a Gaussian point spread function (PSF) with a fitting radius of four pixels and an initial sigma of one pixel. For the 37°C/low calcium condition, the parameters were adjusted where the local maximum instead used 2.0*std (Wave.F1) and a connectivity of 8-neighborhood, and sub-pixel localization used a Gaussian PSF with a fitting radius of three pixels and an initial sigma of one pixel. Once parameter optimization was completed, the custom-made ImageJ macro BatchProcess_ThunderSTORM,ijm was used to iterate ThunderSTORM spot detection over all image files, applying the same parameters for a given biological replicate. Detected spots were linked with cell meshes using the custom MATLAB code analyze_ensemble_spot_intensity.m, detailed further below. Code developed as part of this study has been deposited in the publicly accessible Github code repository (https://github.com/JacobsWagnerLab/published/tree/master/Schubert_et_al_2024). Three independent experiments were performed and analyzed per strain and condition.

Elaborating on the MATLAB script analyze_ensemble_spot_intensity.m, each cell was quantified by averaging whole cell intensity, recording the number of ThunderSTORM-detected spots, and noting the cell length. Only cells with length of at least 1µm were kept to rule out incorrect cell segmentations. Each cell was identified and measured from its original Oufti mesh. There, the fluorescent GFP images were background-subtracted, then cell contours were used to obtain the boundaries of the cell area. Cellular fluorescence was determined by Oufti [93]. To link ThunderSTORM spots, (*x*,*y*) localizations, to Oufti meshes, the cell meshes were first converted to MATLAB polygons using the function *polyshape*. The function *inpolygon* was then used to feed the (*x*,*y*) localizations and polygon shape to find if any spots occurred inside the cell. For Fig 3 C-E, cells with zero spot were excluded from the analysis.

To assess what fraction of cells were devoid of GFP signal, strains were grouped by condition (inducing vs. non-inducing as 37°C/low calcium vs. 26°C, respectively). Each strain in the condition was subsetted by random sampling without replacement to match the lowest N value (7,033 cells in 26°C and 10,003 cells in 37°C/low calcium) using the function *datasample*. These data were then bootstrapped 100 times to recalculate histograms and determine the error for each bin (using the custom function *Hist_Errors*). Since the Δ*pcnB* conditions had clear bimodal distributions for low/high fluorescent states, an Otsu threshold was calculated using the function *otsuthresh* to find the split between them [97]. Cell with intensity values below this threshold were deemed cells without plasmid content. To compare these two distribution shapes statistically, a Kolmogorov-Smirnov (KS) test was used via the function *kstest2*.

Images were visualized using NIS Elements and Fiji. Image pixel intensity values were auto-scaled for display using Fiji, and all images were converted to RGB format for display. Data were plotted using GraphPad Prism 10.2.0, and Adobe Illustrator was used to generate figures.

### *In vitro* pYV plasmid stability assay

Cells were struck for single colonies onto LB plates and grown at 26°C for 1.5-2 days. To test plasmid stability after 26°C-only passage, single colonies were then patched onto LB plates for ∼16 hours at 26°C before heavily streaking for single colonies onto fresh LB plates at 26°C for 1.5-2 days. To test plasmid stability after exposure to T3SS-inducing conditions (37°C), the same colonies were also patched onto LB plates supplemented with 20 mM sodium oxalate to induce T3SS activity for ∼16 hours at 37°C before streaking for single colonies onto LB plates and returning to 26°C to grow for 1.5-2 days. Final plasmid stability was determined for both conditions by patching single colonies from the final 26°C streak on LB plates supplemented with 20 mM sodium oxalate and containing 1% Congo red. The percentage of cells that retained the pYV plasmid was calculated by dividing the number of red (pYV+) CFU by the total number of CFU. Single colony PCR was used on a subset of patches to validate that red CFU retain pYV and while white CFU do not.

### *In vitro* T3SS activity assay

Quantification of secreted *Yersinia* T3SS cargo was carried out as previously described [98]. *Y. pseudotuberculosis* cultures were grown shaking overnight at 26°C before subculturing in LB low calcium media (LB containing 20 mM sodium oxalate and 20 mM MgCl_2_). Cultures were grown shaking at 26°C for 1.5 hours before shifting to 37°C/shaking for an additional 1.5 hours. Cultures were normalized to the same number of cells by normalizing OD_600_ values and pelleted at 13,200 rpm for 15 min at room temperature. Supernatants were transferred to new tubes and proteins precipitated with 10% trichloroacetic acid (TCA). A bovine serum albumin (BSA) protein control was spiked into each sample immediately before the addition of TCA. Samples were incubated in TCA on ice for at least 1h before pelleting at 13,200 rpm for 15 min at 4°C. Pellets were then washed twice with ice-cold 100% acetone and resuspended in final sample buffer (FSB) containing 0.2M dithiothreitol (DTT). Samples were boiled for 10 min at 95°C before loading and separating on a 12.5% SDS-PAGE gel. Gels were stained with Coomassie blue to visualize proteins and imaged using Bio-Rad Image Lab Software Quantity and Analysis tools.

### *Shigella* T3S secretion assay

*S. flexneri* cultures were grown with shaking overnight in tryptic soy broth (TCS) media with 15 µg/mL tetracycline and 50 µg/mL kanamycin at 37°C. In the AM, the cultures were sub-cultured in TCS and grown for 2.5 hours with shaking at 37°C to reach OD_600_ ∼1. Cultures were normalized to the same number of cells based on the OD_600_ and pelleted at 13,000 rpm for 1 min before resuspending in PBS and 10 µM Congo red and incubated for 30 min shaking at 37°C. Pellets and supernatants were separated by centrifugation and supernatants were transferred to new tubes and precipitated with 10% TCA. Supernatant samples were incubated on ice and subsequently pelleted at 15000 rpm for 15 min at 4°C. Pellets were then washed with ice-cold 100% ethanol and resuspended in 1x loading dye buffer with 5% beta-mercaptoethanol. Samples were boiled for 10 min at 95°C before loading on a 12% SDS-PAGE gel for western blot analysis.

### RNA isolation and RNA-seq analysis

*Y. pseudotuberculosis* IP2666pIB1 strains were grown in 3 mL LB overnight at 26°C and diluted the following morning to an OD_600_ of 0.2 in 5 mL fresh LB. Each strain was diluted into two separate cultures (one for each temperature). For 26°C samples, cultures were diluted in plain LB media and grown shaking at 26°C for 3h before extracting RNA. For 37°C samples, cultures were diluted into LB low calcium media (containing 20mM sodium oxalate and 20mM MgCl_2_) and grown shaking at 26°C for 1.5 hours before shifting to 37°C to growing shaking for an additional 1.5 hours. After incubating, 1-2mL of each culture was collected and pelleted for 5 min at 4,000 rpm. Supernatants were removed and cell pellets were resuspended in 1mL RNA Protect (Qiagen) and total RNA was isolated using the RNeasy minikit (Qiagen) according to the manufacturer’s protocol. DNA was removed from samples using the TURBO DNA-free kit (Life Technologies/Thermo Fischer) and RNA quality was determined using an Agilent 2000 Bioanalyzer. Samples were DNase treated with Invitrogen DNase. Library preparation was performed using Illumina’s Stranded Total RNA Prep Ligation with Ribo-Zero Plus kit and 10 bp IDT for Illumina indices. Sequencing was done on a NovaSeq 6000 giving 2×51bp reads.

Demultiplexing, quality control, and adapter trimming was performed with bcl-convert (v4.0.3). Read mapping was performed with HISAT2 [99]. Read quantification was performed using Subread’s featureCounts functionality [100]. Read counts were normalized using edgeR’s Trimmed Mean of M values (TMM) algorithm (Supplementary Dataset 1)[101]. Subsequent values were then converted to counts per million (cpm). Differential expression analysis was performed using edgeR’s exact test for differences between two groups of negative-binomial counts with an estimated dispersion value of 0.1. Genes for which the |logFC| > 1 and p < 0.05 were considered differentially expressed (Supplementary Dataset 1).

Heatmaps for pYV-encoded genes were generated using average TMM values via ClustVis [102]. RNA-Seq data are available for viewing, and the track hub data can be found at http://lowelab.ucsc.edu/hubs/yersinia_pcnB_pil_expression/hub.txt on the UCSC Microbial Genome Browser. For these data, chromosomal reads were normalized separately from pYV-encoded reads due to differences in pYV PCN among samples. For this analysis, raw paired reads were aligned to the *Y. pseudotuberculosis* IP2666pIB1 genome (ASM381434v1) using Bowtie2 (v.2.3.5.1) with default options. Bam files were further processed to remove PCR duplicates using samtools (v1.15.1) as follows, samtools sort -n {input_bam} | samtools fixmate -m - - | samtools sort - | samtools markdup -r - - | samtools view -bq 10 -f 0×1 - > {output_bam}. Finally, bamCoverage-deeptools (v3.5.4) was used to normalize the chromosomal read counts using the following options: --normalizeUsing CPM --ignoreForNormalization NZ_CP032567.1 - -outFileFormat bedgraph --binSize 1 --region NZ_CP032566.1. For the non-normalized pYV reads we used bamCoverage-deeptools with options --ignoreForNormalization NZ_CP032566.1 --outFileFormat bedgraph --binSize 1 --region NZ_CP032567.1. Finally, the separate bedgraph files were concatenated and converted to bigwig format using bedGraphToBigWig (v4) with default options.

### Western blot analysis

Cells were grown at the indicated temperature before preparing samples for western blotting. Samples were normalized to the same number of cells using OD_600_ at the time of sample collection. Bacterial pellets were then resuspended in FSB containing 0.2M DTT and boiled at 95°C for 15 minutes. Secreted proteins were precipitated using TCA in the same way as described for the *in vitro* secretion assay above. The same volume of each sample was separated on a 12.5% SDS-PAGE gel and proteins were transferred to a blotting membrane (Immunobilon-P) using a wet mini trans-blot cell (Bio-Rad) at 4°C. Blots were blocked for one hour in Tris-buffered saline containing Tween 20 and 5% milk. Blots were probed with anti-RpoA (gift from Melanie Marketon), goat anti-YopE (Santa Cruz Biotech), mouse M2 anti-FLAG (Sigma), and horseradish peroxidase-conjugated secondary antibodies (Santa Cruz Biotech). Bands were visualized using a BioRad ChemiDoc and quantification analysis was performed using Image Lab software (Bio-Rad).

For *S. flexneri* strains, samples were separated by a 12% SDS-PAGE gel and transferred to a nitrocellulose membrane. Blots were probed with anti-IpaB, anti-IpaC, and anti-IpaD antibodies, and anti-rabbit secondary antibodies. Bands were visualized using SuperSignal West Pico PLUS Chemiluminescent Substrate on an Azure 300 Imager.

### Mouse Infections

C57BL/6 mice were purchased from Jackson Laboratory. Six-to eight-week-old C57Bl/6 mice were infected with 1-3×10^3^ *Y. pseudotuberculosis* via intraperitoneal (IP) injection as previously described [103]. In brief, LB overnight cultures of *Y. pseudotuberculosis* were grown shaking at 26°C before spinning down at 5000 rpm and resuspending in sterile PBS (pH 7.4). CFU used for the inoculum cultures were confirmed to contain pYV prior to infection using single colony PCR with primers specific to the pYV plasmid. Samples were then diluted in PBS such that 100 µL contained 1-3×10^3^ cells. Inoculums were plated to confirm the inoculation dose and to estimate the percentage of cells that were pYV+ by patching onto LB plates supplemented with 20 mM sodium oxalate and containing 1% Congo red dye. The plasmid retention of the starting inoculums was found to be 100% for wild type and 81% on average for Δ*pcnB*. Mice were injected with 100 µL of inoculum and spleens and livers harvested four days post-inoculation.

Organs were homogenized and serial dilutions of the homogenate plated to determine CFU per gram (CFU/g) of tissue.

### Construction of PAP I expression vector

The *pcnB* ORF was PCR amplified from *Y. pseudotuberculosis* genomic DNA using the primer pair FpcnB_pTrc/RpcnB_pTrc and ligated into pTrc99a previously digested with NcoI and Xbal using NEBuilder HiFi DNA Assembly kit (New England Biolabs, Inc.). Constructs were transformed into DH5α *E. coli* for propagation and sequence verified via Sangar sequencing. The resulting pTrc99a::PAPI recombinant expression vector was introduced to wildtype and Δ*pcnB Y. pseudotuberculosis* by electroporation and selection on LB plates containing carbenicillin. Expression of PAPI was induced by plating onto LB plates containing 0.2 mM IPTG.

### Spot tests to analyze antibiotic resistance conferred by pGFPuv, pTrc99, and pNF06

Wildtype and Δ*pcnB Y. pseudotuberculosis* IP2666pIB1 were electroporated with 1 µL of pGFPuv [Carb^R^] (Gifted by Dr. Manel Camps), pTrc99a, or pNF06::ccdAB [Kan^R^] (Gifted by Dr. Laurence Van Melderen) and single colonies were selected by plating on LB plates containing carbenicillin (pGFPuv, pTrc99a) or kanamycin (pNF06). For spot test dilution series analyses, overnight cultures were grown at 26°C in the presence of antibiotics before diluting in PBS (pH 7.4) and making a series of 1:10 dilutions. Serial dilutions were spotted onto LB plates containing no antibiotic or LB plates containing the selective antibiotic and plates were visualized after 1-1.5 days of growth at 26°C.

Δ*ipa2.5::tet^R^* and Δ*ipa2.5*Δ*pcnB S. flexneri* M90Twere electroporated with pTrc99a, and single colonies were selected by plating on LB plates containing ampicillin. For spot tests, overnight cultures were grown at 30°C in the presence of antibiotics before diluting in PBS (pH 7.4) and making a series of 1:10 dilutions. Serial dilutions were spotted onto LB plates containing no antibiotic or LB plates containing the selective antibiotic and plates were visualized after 1 day of growth at 30°C.

### pGFPuv plasmid loss curve

Wildtype and Δ*pcnB Y. pseudotuberculosis* IP2666pIB1 lacking pYV (pYV-) were electroporated with pGFPuv as described above. Overnight cultures were grown at 26°C in the presence of carbenicillin. Overnight cultures were diluted 1:1000 into LB media without antibiotics to allow loss of the plasmid. To assess plasmid retention at each timepoint, aliquots of the cultures were plated onto LB agar and incubated at 26°C. Fifty single colonies from each sample were patched onto LB agar containing carbenicillin. Percentage of plasmid retention was calculated as the number of patched colonies able to grow in the presence of carbenicillin relative to the total number of patched colonies.

## Acknowledgments

We would like to acknowledge Dr. Manel Camps (UC Santa Cruz) for providing the pGFPuv plasmid as well as general advice and guidance on this work. We would also like to acknowledge Dr. Laurence Van Melderen (Université Libre de Bruxelles) for providing us with the pNF06::empty and pNF06::*ccdAB* constructs. Finally, we would like to thank Dr. Petra Dersch (Universität Münster) for providing general advice and technical support for this project.

## Funding

This study was supported by National Institutes of Health (www.NIH.gov) grant R01AI141511 (to VA), R01AI128360 (to CFL), and the Swedish Research Council 2018-02376 and 2022-00741 (to HW). CJW is an investigator of the Howard Hughes Medical Institute. HNL was supported by NIH K99AI139281. KS was supported by an NIH T32 GM008646 training grant. The funders had no role in study design, data collection and analysis, decision to publish, or preparation of the manuscript.

## Competing interests

The authors have declared that no competing interests exist.

**Figure S1. A transposon insertion into *pil* leads to a constitutively high pYV PCN, elevated T3SS expression, and a severe growth defect under T3SS-inducing conditions. (A)** HEK293T cells expressing an NFκB luciferase reporter were left uninfected (uninf) or were infected with an effectorless Δyop6 strain (Δ*yopHEMOJ*), a Δ*yscNU* T3SS-deficient strain, or *pil::Tn* in the *Y. pseudotuberculosis* IP2666pIB1 background. Averages of three independent replicates ± standard error the mean are shown (Student t-test, p<0.04). **(B)** Schematic representation of the location of *pil*. The *pil* locus is located ∼123bp away from the *copB* gene. **(B)** Wildtype, *pil::Tn*, and Δ*pil* IP2666pIB1 were grown at 37°C/low calcium and secreted T3SS cargo proteins (Yops) were precipitated and visualized by Coomassie blue staining. Bovine serum albumin (BSA) serves as a loading and precipitation control. Data shown is representative of three independent replicates. **(D-E)** Relative pYV PCN was estimated for strains in the YPIII/pIBX background of *Y. pseudotuberculosis* using a luciferase plasmid copy number assay. For each timepoint, luminescence was measured and normalized to cell density (OD_600_). Representative data from three independent experiments are shown. **(E)** Serial dilutions for wildtype YPIII/pIBX and the *pil*::pNQ congenic *Y. pseudotuberculosis* strains were spotted onto low calcium agar plates for ∼16 hours at 37°C before imaging. The yellow arrow indicates an example of a large colony in the *pil*::pNQ mutant background that represents a candidate suppressor mutant.

**Figure S2. T3SS gene mRNA levels are higher in the *pil*::Tn mutant and lower in the Δ*pcnB* mutant compared to wildtype.** Heat maps displaying relative expression of pYV-encoded genes from RNA-seq analysis of *Y. pseudotuberculosis* IP2666pIB1 strains grown at 26°C or 37°C/low calcium.

**Figure S3. The *pil*::Tn strain exhibits normal temperature and calcium regulation of T3SS activity.** Strains were grown in regular LB or low calcium LB media, at either 26°C or at 37°C. Secreted proteins in the supernatant were precipitated and visualized by SDS-PAGE and Coomassie blue staining. Bovine serum albumin (BSA) was added as a loading and precipitation control. Data shown is representative data of three biological replicates.

**Figure S4. Transposon insertion in the *pil* locus alters RNA-Seq read patterns at the *cop*-*rep* locus.** Reads from RNA-seq analysis mapped to the *cop-rep* locus and adjacent *pil* locus from wildtype and *pil*::Tn (*pil*) *Y. pseudotuberculosis* IP2666pIB1 grown at 26°C or 37°C/low calcium. The putative *pil* locus is indicated by a black box and the approximate location of the Tn insertion is indicated with an arrow.

**Figure S5. The *pcnB* gene is required for plasmid-mediated antibiotic resistance in *Y. pseudotuberculosis.*** Wildtype and Δ*pcnB Y. pseudotuberculosis* IP2666pIB1 harboring either pBAD33 or pACYC184 were grown overnight in LB supplemented with 20 µg/mL chloramphenicol, and the overnight cultures spotted onto LB agar lacking or containing chloramphenicol to assess plasmid retention. Plates were incubated at 26°C for ∼16 hours before imaging. Each experiment was carried out for a total of two independent experiments.

**Figure S6. *Y. pseudotuberculosis* isolated from infected livers demonstrate similar plasmid loss and T3SS activity phenotypes as their parental strains.** Independent colony forming units from infected mouse livers were isolated and assessed for their ability to retain pYV and carry out T3SS activity. WT-3 and WT-5 are wildtype IP2666pIB1-derived isolates from two different mice. Δ*pcnB*-2-9, Δ*pcnB*-2-10, Δ*pcnB*-3-1, and Δ*pcnB*-3-2 are Δ*pcnB*-derived isolates from two different mice. **(A)** Liver isolates and their parental strains were incubated overnight at 26°C in LB or 37°C in LB/low calcium media. Colonies were patched onto low calcium media containing Congo red to induce T3SS activity and assess the percentage of colonies that retained pYV (i.e.-red colonies). **(B)** Liver isolates and their parental strains were incubated in LB low calcium media at 37°C, and secreted proteins visualized by Coomassie blue staining. Bovine serum albumin (BSA) was added as a loading and precipitation control. Data shown is representative of two independent experiments.

**Figure S7. Loss of PAP I does not affect *Y. pseudotuberculosis* growth following stress exposure. (A)** Wildtype and Δ*pcnB Y. pseudotuberculosis* IP2666pIB1 cultures were grown to mid-log phase at 26°C before shifting to 42°C for 1 hour to induce heat shock. Spot dilutions were plated onto LB plates and incubated at 26°C for ∼16 hours before imaging. **(B)** Strains in the IP2666pIB1 background were grown at 26°C overnight (stationary phase) before diluting and spotting into LB plates. Plates were incubated at 26°C for 4 hours before transferring to 4°C and allowing growth for ∼2 days prior to imaging. **(C)** Wildtype and Δ*pcnB Y. pseudotuberculosis* IP2666pIB1 were grown at 37°C (low calcium) to mid-log phase before inducing stress. Growth was assayed by plating spot dilutions following stress exposure. Stress was induced by adjusting media to pH=3.0 to induce acid shock or by addition of 800Mm NaCl_2_ to induce osmotic shock.

**Figure S8. PAP I from *Y. pseudotuberculosis* and *Y. pestis* are 100% conserved.** Predicted PAP I sequences from *Y. pseudotuberculosis* IP2666pIB1 and *Y. pestis* KIM5.

**Figure S9. *Y. pseudotuberculosis* displays normal T3SS activity in EZ-RDM low calcium media.** *Y. pseudotuberculosis* IP2666pIB1 wildtype, Δ*pcnB*, and pYV-strains were grown in either low calcium LB or low calcium EZ-RDM media. The pYV-strain cured of pYV was used as a negative control [103]. Secreted proteins were visualized by Coomassie blue staining. Bovine serum albumin (BSA) was added as a loading and precipitation control. Data shown is representative of two independent experiments.

**Table S1. Phenotypic categories and mutations identified in suppressor screen.**

**Table S2. *Y. pseudotuberculosis* strains used in this study.**

**Table S3. Primers used in this study.**

**Table S4. Plasmids used in this study.**

**Dataset S1. Transcriptome analysis of *Y. pseudotuberculosis* IP2666pIB1, *pil*::Tn, Δ*pcnB*, and *pil*::Tn/Δ*pcnB* congenic strains.**

## Notes

### Competing Interest Statement

The authors have declared no competing interest.

### Summary of Updates

We have added supplemental data figures.

