## Supplemental Figures for "The polyadenylase PAPI is required for virulence plasmid maintenance in pathogenic bacteria"

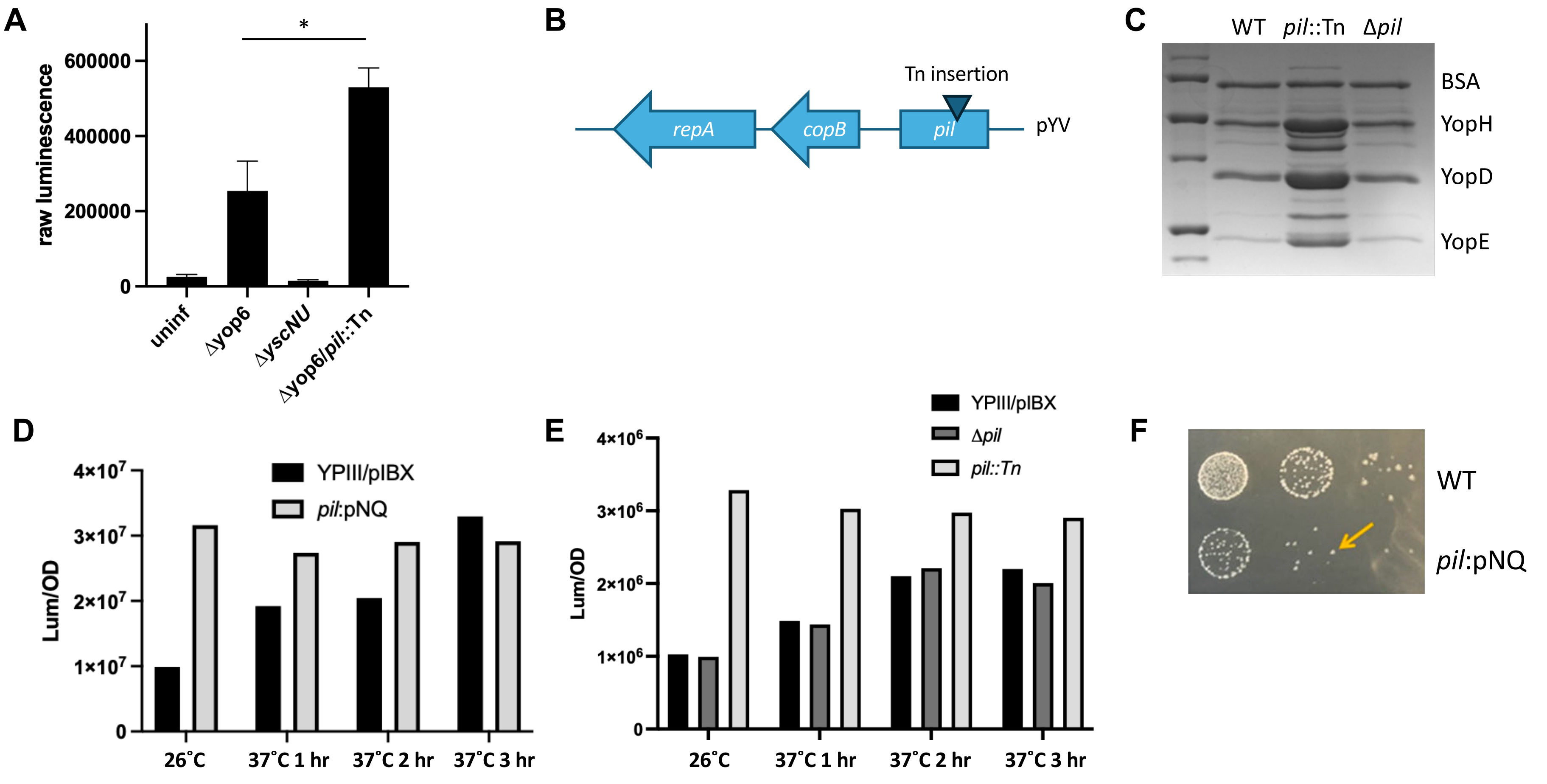
A
raw luminescence
600000
400000
200000
0
*
uninf
Δyop6
ΔyscNU
Δyop6/pil::Tn
B
Tn insertion
repA
copB
pil
pYV
C
WT
pil::Tn
Δpil
BSA
YopH
YopD
YopE
D
Lum/OD
4×10^7
3×10^7
2×10^7
1×10^7
0
YPIII/pIBX
pil:pNQ
26˚C
37˚C 1 hr
37˚C 2 hr
37˚C 3 hr
E
Lum/OD
4×10^6
3×10^6
2×10^6
1×10^6
0
YPIII/pIBX
Δpil
pil::Tn
26˚C
37˚C 1 hr
37˚C 2 hr
37˚C 3 hr
F
WT
pil:pNQ

Figure S1

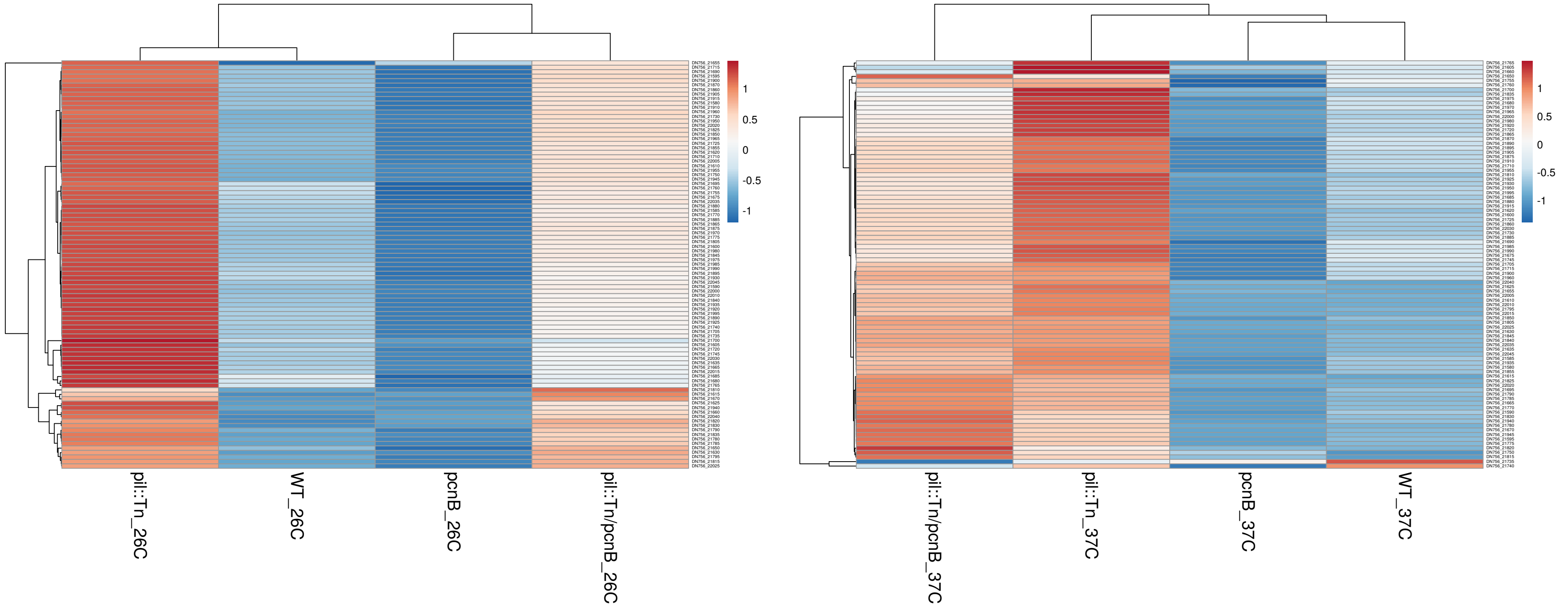

1
0.5
0
-0.5
-1
pil::Tn_26C
WT_26C
pcnB_26C
pil::Tn/pcnB_26C
pil::Tn/pcnB_37C
pil::Tn_37C
pcnB_37C
WT_37C


Figure S2

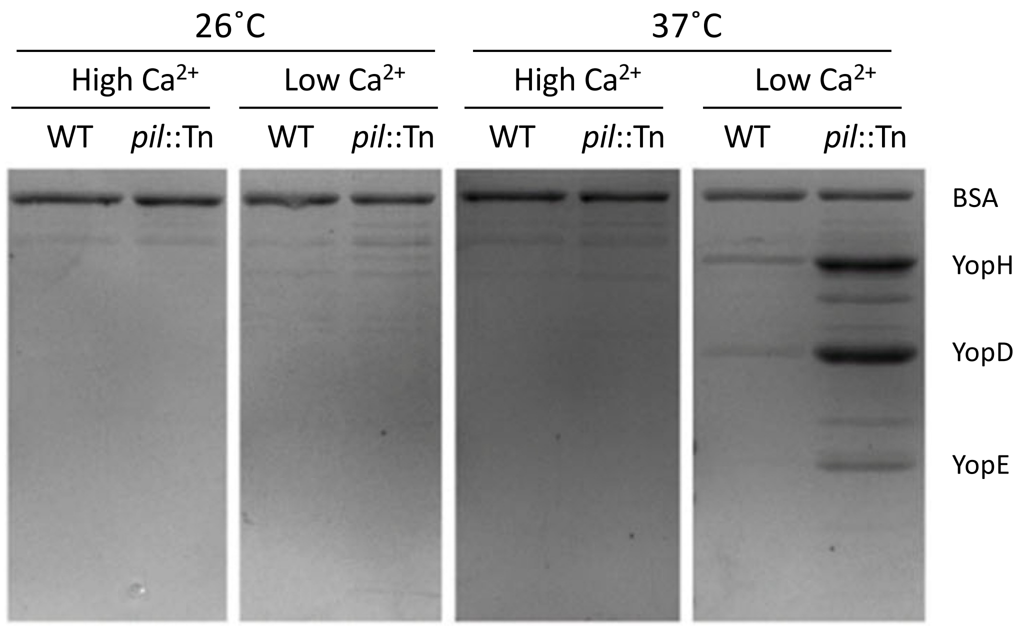
26˚C
37˚C
High Ca2+
Low Ca2+
High Ca2+
Low Ca2+
WT
pil::Tn
WT
pil::Tn
WT
pil::Tn
WT
pil::Tn
BSA
YopH
YopD
YopE

Figure S3

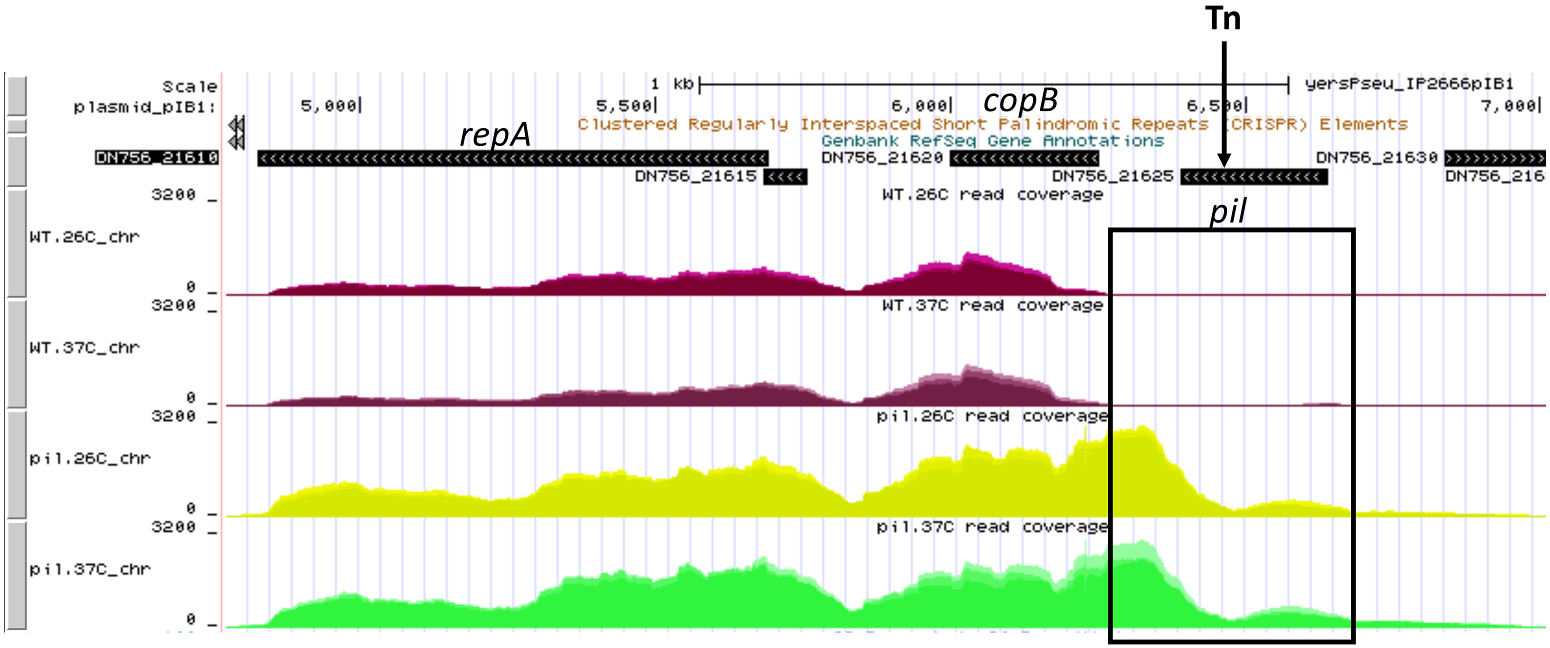

Tn
Scale
plasmid_pIB1:
1 kb
yersPseu_IP2666pIB1
5,000
5,500
6,000
6,500
7,000
copB
repA
Clustered Regularly Interspaced Short Palindromic Repeats (CRISPR) Elements
Genbank RefSeq Gene Annotations
DN756_21610
DN756_21615
DN756_21620
DN756_21625
DN756_21630
DN756_216
pil
WT.26C read coverage
WT.37C read coverage
pil.26C read coverage
pil.37C read coverage
WT.26C_chr
WT.37C_chr
pil.26C_chr
pil.37C_chr
3200
0


Figure S4

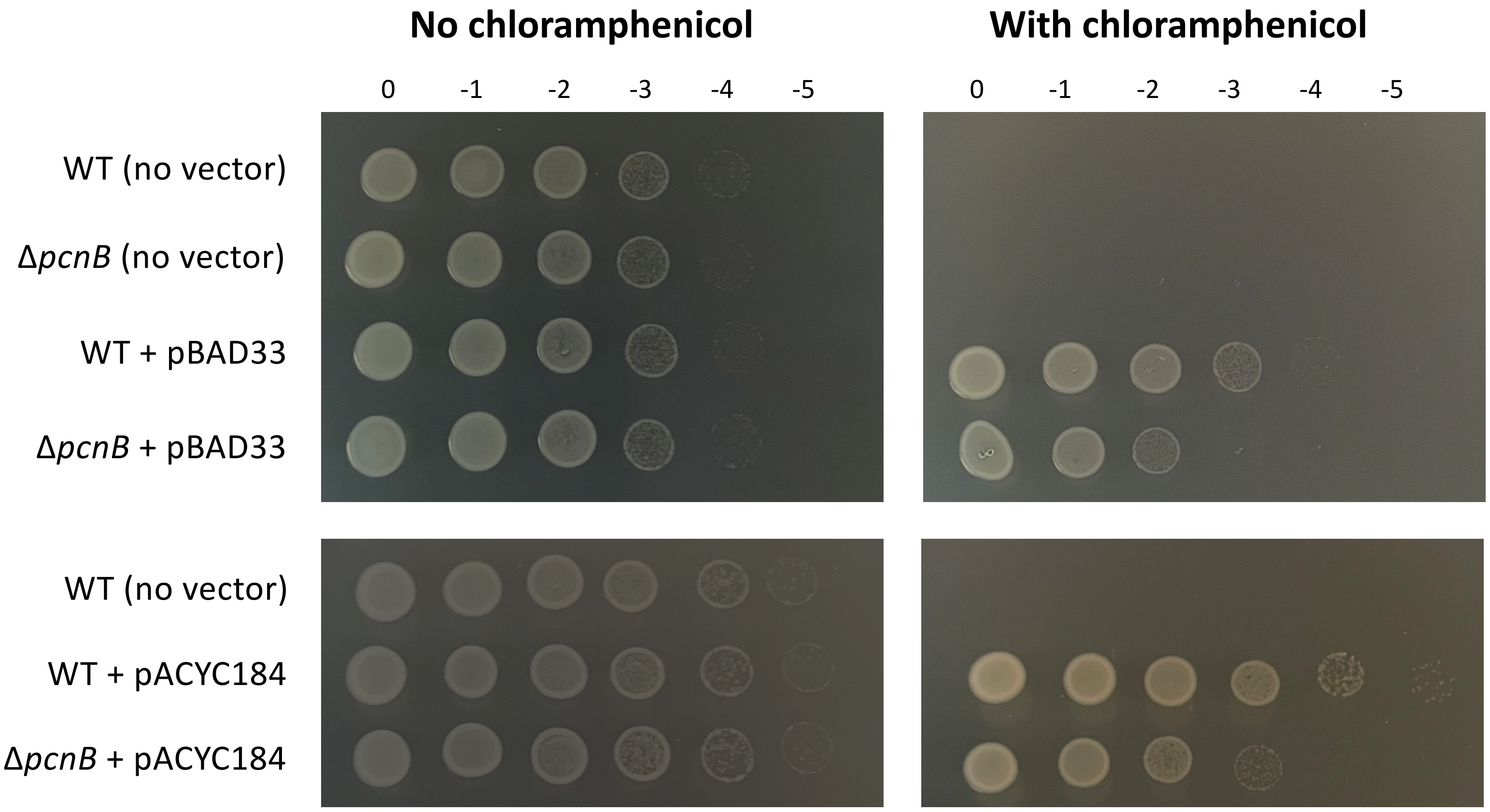
No chloramphenicol
With chloramphenicol
0
-1
-2
-3
-4
-5
0
-1
-2
-3
-4
-5
WT (no vector)
ΔpcnB (no vector)
WT + pBAD33
ΔpcnB + pBAD33
WT (no vector)
WT + pACYC184
ΔpcnB + pACYC184

Figure S5

**A**

| | **26˚C** | **37˚C** |
|---|---|---|
| WT | 100% | 94% |
| Δ*pcnB* | 89% | 39% |
| WT-3 | 100% | 94% |
| WT-5 | 100% | 67% |
| Δ*pcnB*-2-9 | 85% | 16% |
| Δ*pcnB*-2-10 | 92% | 25% |
| Δ*pcnB*-3-1 | 83% | 8% |
| Δ*pcnB*-3-2 | 91% | 23% |

**B**

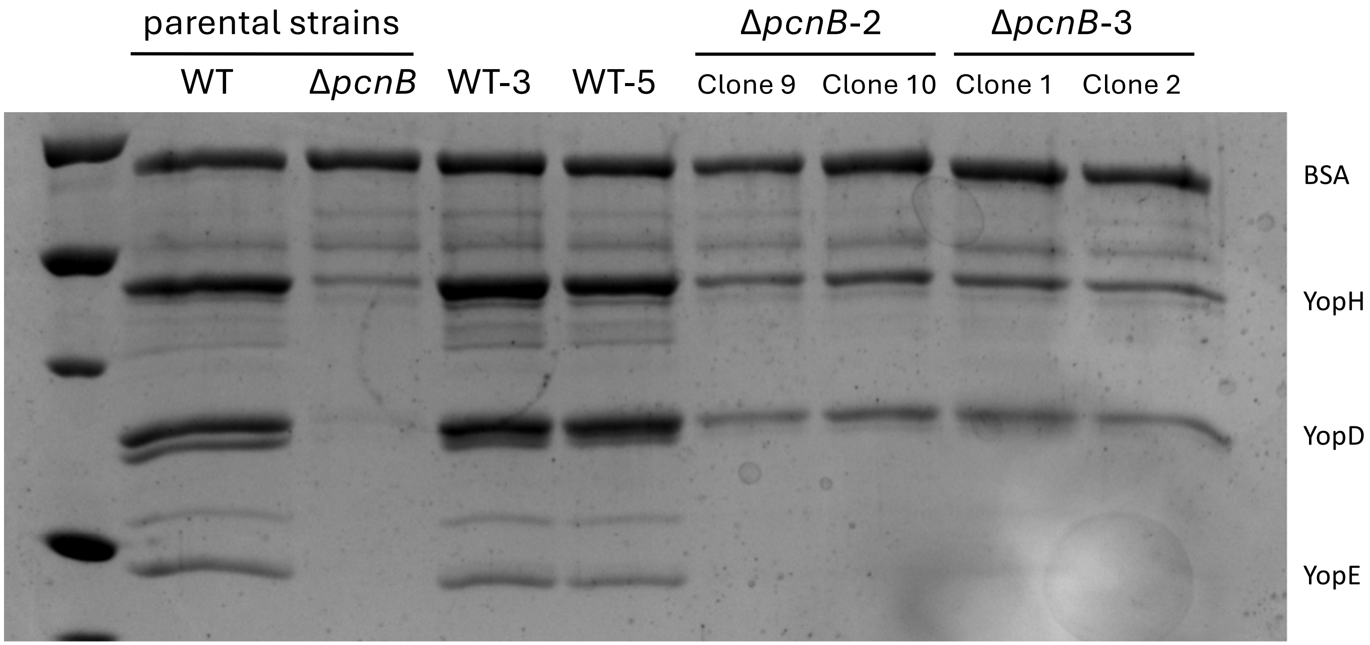


Figure S6

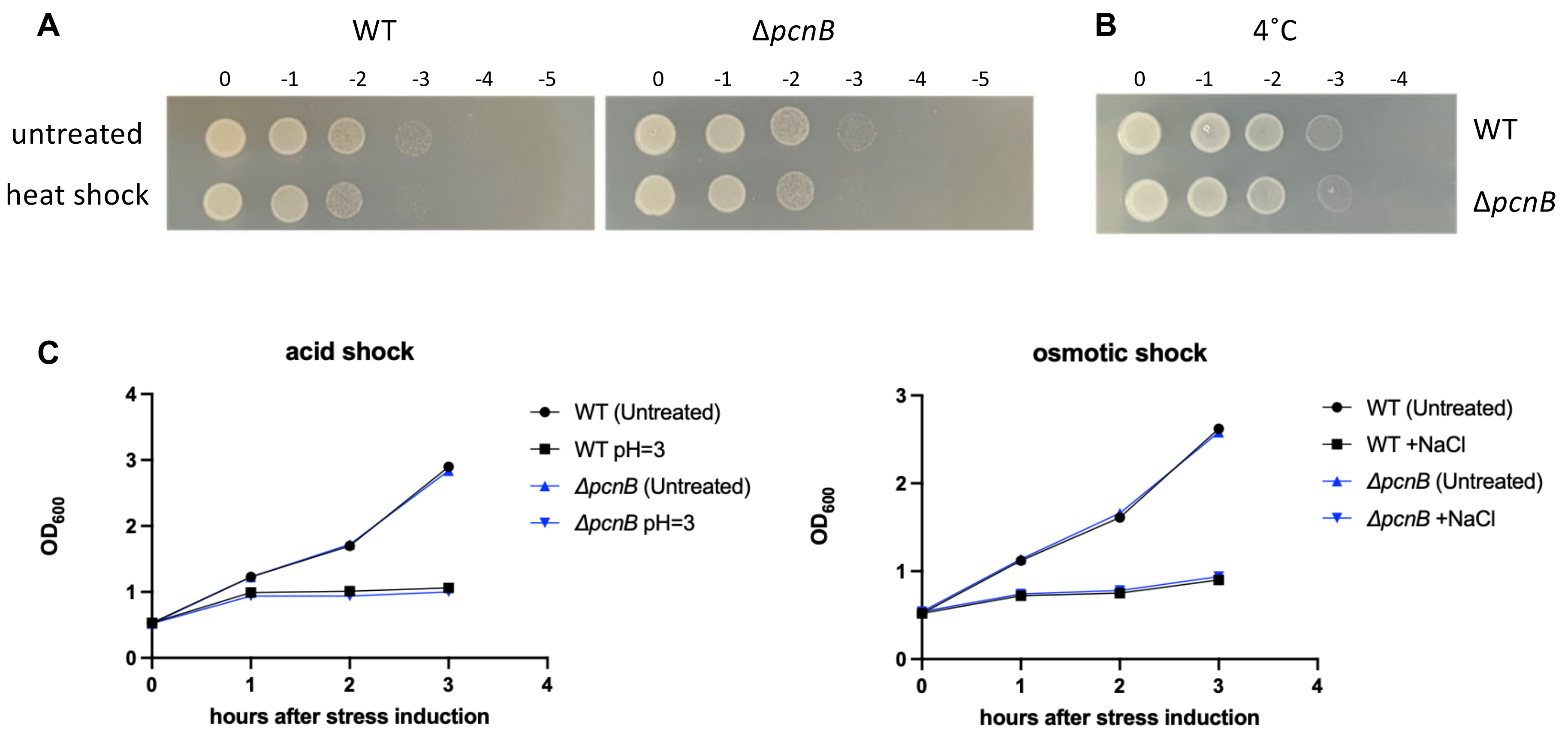
A
WT
ΔpcnB
0 -1 -2 -3 -4 -5
0 -1 -2 -3 -4 -5
untreated
heat shock
B
4˚C
0 -1 -2 -3 -4
WT
ΔpcnB
C
acid shock
OD600
hours after stress induction
WT (Untreated)
WT pH=3
ΔpcnB (Untreated)
ΔpcnB pH=3
osmotic shock
OD600
hours after stress induction
WT (Untreated)
WT +NaCl
ΔpcnB (Untreated)
ΔpcnB +NaCl

Figure S7

```
Y.pseudotuberculosis(IP2666)      MFTRVANFCRKVLIREDKTVRDDKARKDKVPGEDNVARKERRPARAHTGRKGHAVSSSEQ  60
Y.pestis(Kim5)                    MFTRVANFCRKVLIREDKTVRDDKARKDKVPGEDNVARKERRPARAHTGRKGHAVSSSEQ  60
                                  ************************************************************

Y.pseudotuberculosis(IP2666)      RQMAIIPRDQHNISRRDISDNALKVLYRLNKSGYEAYLVGGGVRDLLLGRKPKDFDITTS  120
Y.pestis(Kim5)                    RQMAIIPRDQHNISRRDISDNALKVLYRLNKSGYEAYLVGGGVRDLLLGRKPKDFDITTS  120
                                  ************************************************************

Y.pseudotuberculosis(IP2666)      ATPEQVRKLFRNCRLVGRRFRLAHVMFGPEIIEVATFRGHHEQQQAEDSDKNSSQQAQNG  180
Y.pestis(Kim5)                    ATPEQVRKLFRNCRLVGRRFRLAHVMFGPEIIEVATFRGHHEQQQAEDSDKNSSQQAQNG  180
                                  ************************************************************

Y.pseudotuberculosis(IP2666)      MLLRDNIFGSIEDDAQRRDFTINSLYYGISDFALRDYTGGLRDLKEGIIRLIGDPETRYR  240
Y.pestis(Kim5)                    MLLRDNIFGSIEDDAQRRDFTINSLYYGISDFALRDYTGGLRDLKEGIIRLIGDPETRYR  240
                                  ************************************************************

Y.pseudotuberculosis(IP2666)      EDPVRMLRAVRFAAKLDMSISPETAEPIPRLASLLREIPPARLFEESLKLLQSGYGYKTY  300
Y.pestis(Kim5)                    EDPVRMLRAVRFAAKLDMSISPETAEPIPRLASLLREIPPARLFEESLKLLQSGYGYKTY  300
                                  ************************************************************

Y.pseudotuberculosis(IP2666)      LKLCEYQLFQPLFPLIARNFTEQHDSPMERILVQVLKNTDHRLHNDQRVNPAFLFAAMLW  360
Y.pestis(Kim5)                    LKLCEYQLFQPLFPLIARNFTEQHDSPMERILVQVLKNTDHRLHNDQRVNPAFLFAAMLW  360
                                  ************************************************************

Y.pseudotuberculosis(IP2666)      YPLIEHAQKLTQESGLAYYDAFALAMNDVLDEECRSLAIPKRITSLVRDIWLLQLRLSRR  420
Y.pestis(Kim5)                    YPLIEHAQKLTQESGLAYYDAFALAMNDVLDEECRSLAIPKRITSLVRDIWLLQLRLSRR  420
                                  ************************************************************

Y.pseudotuberculosis(IP2666)      QGKRAHKLMEHPKFRAAYDLLLLRAEVEKNHELQRLAQWWGEFQEATPTQQKSMLNTLGA  480
Y.pestis(Kim5)                    QGKRAHKLMEHPKFRAAYDLLLLRAEVEKNHELQRLAQWWGEFQEATPTQQKSMLNTLGA  480
                                  ************************************************************

Y.pseudotuberculosis(IP2666)      DPAPRRSRPRRPRKPVPRKEGV        502
Y.pestis(Kim5)                    DPAPRRSRPRRPRKPVPRKEGV        502
                                  **********************
```

Figure S8

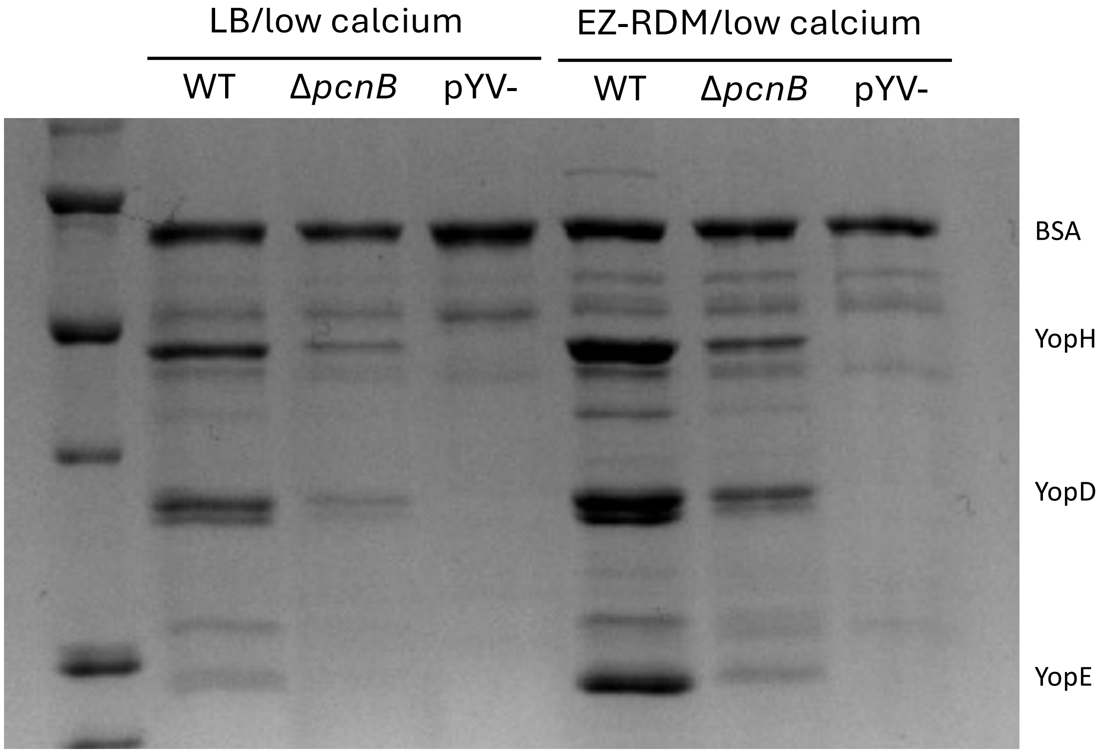

LB/low calcium
EZ-RDM/low calcium
WT
ΔpcnB
pYV-
WT
ΔpcnB
pYV-
BSA
YopH
YopD
YopE


Figure S9
