## Supplemental Tables for "The polyadenylase PAPI is required for virulence plasmid maintenance in pathogenic bacteria"

**Table S1. Phenotypic categories and mutations identified in suppressor screen**

| Strain: | PCN 26°C^1^ | PCN 37°C^1^ | T3SS activity^3^ | Identified Mutation: | Phenotypes: |
| --- | --- | --- | --- | --- | --- |
| pil::pNQ  (parental) | High^2^ | High ^2^ | High ^2^ | N/A |  |
| IS0002 | No Change | No change | No Activity | TGA 🡪 GGA mutation abolishing YscI stop codon | Predicted to decrease or abolish translation of YscI |
| IS0003 | No Change | Lower | Lower | G 🡪 C mutation in tRNA-Gly | Unknown how this mutation impacts pYV PCN and/or T3SS activity. |
| IS0004 | No Change | Lower | Lower | G 🡪 A mutation in *prepA* -35 region | Predicted to decrease  p*repA* promoter firing |
| IS0006 | Lower | Lower | Lower | L291R mutation in PAP I | Mutation lowers PAP I protein levels at 37°C |
| IS0007 | No Change | No Change | No Activity | K175E mutation abolishing the essential lysine in walker box A of YscN | Mutation predicted to impair YscN ATP binding and hydrolysis |
| IS0008 | Lower | Lower | Lower | L291R mutation in PAP I | Mutation lowers PAP I protein levels at 37°C |
| IS0016 | Lower | Lower | Lower | A272E in RNA Pol alpha factor | Mutation may impact RNA Pol binding to some promoters [1] |

^1^ Relative pYV PCN was determined using a luciferase PCN assay with luminescence
 normalized to OD_600_.

^2^ Changes in pYV PCN and T3SS activity relative to the parental strain. The parental strain
 for *pil::pNQ* is wildtype *Y. pseudotuberculosis* YPIII/pIBX and the parental strain for all
 suppressor isolates shown is *pil::pNQ* YPIII/pIBX.

^3^ T3SS activity was determined via a secretion assay of cells grown at 37°C in low calcium
 media.

**Table S2. *Y. pseudotuberculosis* strains used in this study**

| Strain | Background | Mutation(s) | Ref |
| --- | --- | --- | --- |
| Wildtype (WT) | IP2666pIB1 | Naturally lacks full-length YopT | [2] |
| *pil::Tn* | IP2666pIB1 | Tn5 insertion in *pil,* wildtype background | This work |
| *∆pcnB* | IP2666pIB1 | *∆pcnB* | This work |
| *pil::Tn^∆pcnB^* | IP2666pIB1 | *∆pcnB, pil::Tn* background | This work |
| PAP I^His^ | IP2666pIB1 | PAP I with C-6xHis tag | This work |
| PAP I^D2A^ | IP2666pIB1 | D114A+D116A mutations in PAP I (C-6xHis) | This work |
| PAP I^L291R^ | IP2666pIB1 | L291R mutation in PAP I (C-6xHis) | This work |
| PAP I^L291A^ | IP2666pIB1 | L291A mutation in PAP I (C-6xHis) | This work |
| PAP I^FLAG^ | IP2666pIB1 | PAP I with C-3xFLAG tag | This work |
| PAP I^D2A-FLAG^ | IP2666pIB1 | D114A+D116A mutations in PAPI(C-3xFLAG) | This work |
| PAP I^L291R-FLAG^ | IP2666pIB1 | L291R mutation in PAP I (C-3xFLAG) | This work |
| PAP I^L291A-FLAG^ | IP2666pIB1 | L291A mutation in PAP I (C-3xFLAG) | This work |
| *ΔyopHEMOJ* | IP2666pIB1 | *ΔyopHEMOJ* | [3] |
| *ΔyscNU* | IP2666pIB1 | *ΔyscNU* | [4] |
| pYV- | IP2666 | Cured of pYV | [3] |
| *pil::Tn* | IP2666pIB1 | Tn5 insertion in *pil, ∆yopEMOJ* background | This work |
| Wildtype (WT) | YPIII pIBX | Naturally lacks full-length YopT; pIBX is pYV encoding two copies of the *luxCDABE* operon and a kanamycin resistance gene | [5] |
| *pil::pNQ* | YPIII pIBX | pNQ insertion in *pil* | This work |
| *∆pcnB* | YPIII pIBX | *∆pcnB* | This work |
| ParB-GFP | YPIII pIBX | ParB-C-msfGFP | This work |
| *∆pcnB*^ParB-GFP^ | YPIII pIBX | *∆pcnB* ParB-C-msfGFP | This work |
| PAP I^His^ | YPIII pIBX | PAP I with C-6xHis tag | This work |
| PAP I^D2A^ | YPIII pIBX | D114A+D116A mutations in PAP I (C-6xHis) | This work |
| PAP I^L291R^ | YPIII pIBX | L291R mutation in PAP I (C-6xHis) | This work |
| PAP I^L291A^ | YPIII pIBX | L291A mutation in PAP I (C-6xHis) | This work |
| ∆*ipaH2.5::tet^R^* | M90T | ∆*ipaH2.5* | [6] |
| ∆*ipaH2.5*∆*pcnB* | M90T | ∆*ipaH2.5*∆*pcnB* | This work |
| VP- | M90T | BS176 | [7] |

**Table S3. Primers used in this study**

| **Name** | **Primer Sequence** | **Ref** |
| --- | --- | --- |
| **Primers used to make pCVD442::*∆pcnB*** | |  |
| FpcnB_500up_pCVD | caacataaaggtgaatcccatatgAGAAGATTTTATTATCCGTCG | This work |
| RpcnB_500up_pCVD | ggtaaaattaAATGGTACACCTCGATAG | This work |
| FpcnB_500d_pCVD | gtgtaccattTAATTTTACCATGATCCGGGTC | This work |
| RpcnB_500d_pCVD | acctggcacggctgggacggaagtcTGAGCCGGCGATATTACC | This work |
| **Primers used to make pCVD442::PAPI^His^** | |  |
| FpcnB_pET28 | agtggtggtggtggtggtgctcgagTACCCCTTCTTTACGGGG | This work |
| RpcnB_pET28 | actttaagaaggagatataccatggATTTTTACCCGAGTAGCC | This work |
| FpcnB_500up_His | gaggtgtaccATTTTTACCCGAGTAGCCAATTTC | This work |
| RpcnB_500up_His | catggtaaaaTCAGTGGTGGTGGTGGTG | This work |
| FpET_pB_His_300d | caacataaaggtgaatcccatatgAGGGGAATAAGCTCTCCAAG | This work |
| RpET_pB_His_300d | gggtaaaaatGGTACACCTCGATAGTGG | This work |
| FpcnB_300d_pCVD | ccaccactgaTTTTACCATGATCCGGGTC | This work |
| RpcnB_300d_pCVD | tgacagtctccggaagacggTGAGCCGGCGATATTACC | This work |
| **Primers used to make pCVD442::PAPI^FLAG^** | |  |
| FpcnB_FLAG | caacataaaggtgaatcccaTGAAAAATACCGACCACC | This work |
| RpcnB_FLAG | ctttgtagtcTACCCCTTCTTTACGGGG | This work |
| FpET_FLAG_pcnB | agaaggggtaGACTACAAAGACCATGACG | This work |
| RpET_FLAG_pcnB | ggtaaaattaCATATGGTACCAGCTGCAG | This work |
| FpcnB_500d_pCVD | gtaccatatgTAATTTTACCATGATCCGGGTC | This work |
| RpcnB_500d_pCVD | tgacagtctccggaagacggTGAGCCGGCGATATTACC | This work |
| **Primers used to make pCVD442::ParB-msfGFP** | |  |
| FparB_GFP | caacataaaggtgaatcccatatgTTGCTAATGAGTATCGTC | This work |
| RparB_GFP | cgccttttgaCAAAGAATGTTCCTTTGC | This work |
| FpKD_GFP_parB | acattctttgTCAAAAGGCGAAGAACTTTTTAC | This work |
| RpKD_GFP_parB | tattcaggcaTTATTTATACAATTCATCCATTCCATGAGTGAT TCCTGCCGCAGTGACAAATTC | This work |
| FparB_500d_pCVD | gtataaataaTGCCTGAATAAGATCAGAAC | This work |
| RparB_500d_pCVD | tgacagtctccggaagacggAATCTCCCTAAAGCTATCAC | This work |
| **Q5 mutagenesis primers used to introduce L291R mutation** | |  |
| FpB_L291R_pCVD | ACTCAAGCTGaggCAATCCGGCTAC | This work |
| RpB_L291R_pCVD | GACTCCTCAAACAGGCGG | This work |
| **Q5 mutagenesis primers used to introduce L291A mutation** | |  |
| FpB_L291A_pCVD | ACTCAAGCTGgcgCAATCCGGCTAC | This work |
| RpB_L291A_pCVD | GACTCCTCAAACAGGCGG | This work |
| **Q5 mutagenesis primers used to introduce D114A + D116A (D2A) mutations** | | |
| FpB_D2A_pCVD | tcgctATCACCACCAGCGC | This work |
| RpB_D2A_pCVD | aagcTTTGGGTTTTCTGCCC | This work |
| **Primers used to make pTrc99::PAP I** | | |
| FpcnB_pTrc | atttcacacaggaaacagaccatggATTTTTACCCGAGTAGCCAATTTC | This work |
| RpcnB_pTrc | tgcatgcctgcaggtcgactctagaCAGCGCGATATAGACCCG | This work |
| **Primers used for ddPCR** | | |
| F_Chrom_ddPCR | CCTCACCGATACCGAACGAG | [8] |
| R_Chrom_ddPCR | GTCAGCAGGATAGGGCTACC | [8] |
| F_pYV_ddPCR | CTCTTTGACCTCGGCTTGAG | [8] |
| R_pYV_ddPCR | CGCAGCCGTTAGGACAAATG | [8] |
| **Primers used to make ∆*ipaH2.5*∆*pcnB Shigella*** | | |
| pcnb_F | ggcagaagcacactggcagg | This work |
| pcnb_R | gtggtccccagcgttcagc | This work |
| pcnB_R_op | gacttcacgcaacgtctcccc | This work |
| K2_wanner | cggtgccctgaatgaactgc | [9] |

**Table S4. Plasmids used in this study**

| **Vector/Plasmid** | **Insert** | **Ref** |
| --- | --- | --- |
| pCVD442::empty | None [SacB^+^] | [10] |
| pET28::empty | -C-3xFLAG and -C-6xHis | [11] |
| pKD13-msfGFP | -C-msfGFP | ^[12]^ |
| pCVD442:: *∆pcnB* | *∆pcnB* | This work |
| pCVD442::PAP I^His^ | PAP I-C-6xHis | This work |
| pCVD442::PAP I^L291R^ | PAP I^L291R-C-6xHis^ | This work |
| pCVD442::PAP I^L291A^ | PAP I^L291A-C-6xHis^ | This work |
| pCVD442::PAP I^D2A^ | PAP I^D2A-C-6xHis^ | This work |
| pCVD442::PAP I^FLAG^ | PAP I-C-3xFLAG | This work |
| pCVD442::PAP I^L291R-FLAG^ | PAP I^L291R-C-3xFLAG^ | This work |
| pCVD442::PAP I^L291A-FLAG^ | PAP I^L291A-C-3xFLAG^ | This work |
| pCVD442::PAP I^D2A-FLAG^ | PAP I^D2A-C-3xFLAG^ | This work |
| pCVD442::ParB-msfGFP | ParB-C-msfGFP | This work |
| pTrc99A::empty | None [IPTG-inducible T7] | [13] |
| pTrc99::PAP I | PAP I | This work |
| pGFP-uv | GFPuv | [14] |
| pNF06::ccdAB | ccdAB | [15] |
